# Lung epithelial cell-derived C3 protects against pneumonia-induced lung injury

**DOI:** 10.1101/2022.02.03.478963

**Authors:** Ayşe N. Ozantürk, Sanjaya K. Sahu, Devesha H. Kulkarni, Lina Ma, Ruteja A Barve, Ja’Nia McPhatter, Lorena Garnica, Linus Dannull, Jeremy Kunen, Xiaobo Wu, Steven L. Brody, John P. Atkinson, Hrishikesh S. Kulkarni

## Abstract

The complement component C3 is a fundamental plasma protein for host defense. However, recent work has demonstrated the critical importance of local C3 expression in cell survival. Here we analyzed the effects of local versus peripheral sources of C3 expression in a model of bacterial pneumonia. While mice with global C3 deficiency had severe pneumonia-induced lung injury, those deficient in liver-deficient C3 remain protected, comparable to wildtype mice.

Human lung transcriptome analysis showed secretory epithelial cells are a major source of C3. Mice with a C3 gene ablation from lung epithelial cells had worse pulmonary injury compared to wild type, despite maintaining normal circulating C3 levels. Finally, in human cellular and mouse pneumonia models, we show that C3 reduces epithelial cell death mediated through the alternative pathway component Factor B. Thus, our findings suggest that a locally-derived C3-Factor B pathway protects the lung mucosal barrier.

**One Sentence Summary:** Lung-derived C3 mitigates severe bacterial pneumonia suggesting a novel cytoprotective role at mucosal barrier surfaces independent of circulating C3.

## INTRODUCTION

Pathogens infect and injure mucosal surfaces resulting in a cascade of events including inflammation, tissue damage and systemic dissemination (*1, 2*). In the lung, this process results in severe pneumonia, a leading cause of death, even before the SARS-CoV-2 pandemic (*3, 4*). The outcome in pneumonia is determined by a balance between immune resistance and tissue resilience of the host (*1, 5*). Immune resistance is the host’s ability to eradicate pathogens, whereas tissue resilience is the ability of the lung to withstand the deleterious consequences of infection and inflammation (*1, 5*). The complement system is a family of immune proteins that facilitate immune resistance by attacking microbes to decrease pathogen burden (*6, 7*).

Consequently, genetic and acquired deficiencies of complement characteristically lead to recurrent bacterial infections in both children and adults (*8, 9*). However, there has been an increasing appreciation recently of how the system functions to promote tissue resilience in the setting of acute infections (*10, 11*).

A majority of the complement proteins are primarily liver-derived (*12*). One such protein, C3, is a central component of the complement system, as multiple pathways converge on its activation to C3a and C3b, thereby amplifying the cascade (*13*). The plasma concentration of C3 is approximately 1-2 mg/ml in humans, resulting it in being one of the most common proteins in the circulation after other liver-derived proteins such as albumin and α-2-macroglobulin (*14*). Thus, the protective effects of C3 have largely been attributed to it being sourced from the liver and present in a high concentration in the circulation. However, we and others have previously demonstrated that C3 is also produced outside of the liver by multiple immune and non-immune cell populations (*15–19*). This observation has implications, especially for organs that constantly interact with pathogens and environmental toxins, such as the mucosal surface of the lung (*6, 20*). For example, we have demonstrated that C3 synthesized as well as internalized by lung epithelial cells protects against stress-induced cell death (*16, 21*). Others have demonstrated that chronic C3 upregulation in lung epithelial cells can perpetuate inflammation in SARS-CoV-2 infection (*19*).

It is unknown whether cell-intrinsic C3 production is relevant in vivo. It is also unclear if the protective effects of C3 are purely a result of enhancing immune resistance via its bactericidal activity, or it also contributes to tissue resilience by reducing acute lung injury (ALI). To address these questions, we utilized a model of acute *Pseudomonas aeruginosa (Pa)* pneumonia in mice globally deficient in C3 and report that C3 not only promotes immune resistance but also enhances tissue resilience in vivo by mitigating ALI. Using publicly available human lung datasets and a C3 reporter mouse model, we establish that multiple structural cell types in the lung produce and upregulate C3 in the setting of *Pa* infection. Building on these findings, we generated novel mouse strains and show that lung-derived C3 is functioning locally, independent of circulating (i.e., liver-derived) C3 and is necessary for reducing *Pa*-induced ALI. Using both human in vitro and mouse in vivo infection models, we show that cell-intrinsic Factor B — a component of the alternative pathway and a ligand for C3 — is a key component of tissue resilience, enhancing the ability of the lung to withstand the deleterious consequences of infection and inflammation. Our findings potentially refine the approach for therapeutically targeting the complement system in the lung with the goal of mitigating bacterial pneumonia severity.

## RESULTS

### C3 protects against severe bacterial pneumonia

C3-deficient mice had worse histological evidence of lung injury when compared to wildtype (WT) mice at 24 h after *Pa* infection (**Fig 1A**). They had worsened physiological function, as demonstrated by reduced dynamic compliance compared to WT mice at 24 h post-infection (**Fig 1B**), as well as a higher airway resistance (**Fig S1A**), both measured by whole body plethysmography. They also had elevated alveolar capillary-barrier permeability as demonstrated using the wet-dry ratio (**Fig S1B**). Consistent with prior reports, male C3-deficient mice had a significantly worse injury relative to female C3-deficient mice (**Fig. S1C**) (*22*). Hence, C3-deficient male mice were used for subsequent experiments. We next analyzed inflammatory gene expression. C3-deficient mice had an exaggerated inflammatory response, as determined by increased lung *Ccl2* and *Il1b* expression (**Fig 1C**), the proportion of bronchoalveolar lavage (BAL) Ly6C^hi^ monocytes (**Fig S1D**) and BAL CCL2 and IL-1β levels (**Fig S1E and 1F**). The BAL levels of IL-6 and TNF-α showed a trend towards significance in C3-deficient mice compared to WT mice (**Fig S1G and S1H**). Of note, infected C3-deficient mice had lower expression of epithelial cell genes, such as *Scgb1a1* (representing secretory airway epithelial cells) and *Sftpc* (representing alveolar epithelial cells) compared to infected WT, suggesting C3 protects against epithelial cell loss (**Fig 1C**). However, C3-deficient mice also had higher lung (**Fig 1D**) and splenic bacterial counts (**Fig 1E**) at 24 h post-*Pa* infection.

**Figure 1.**
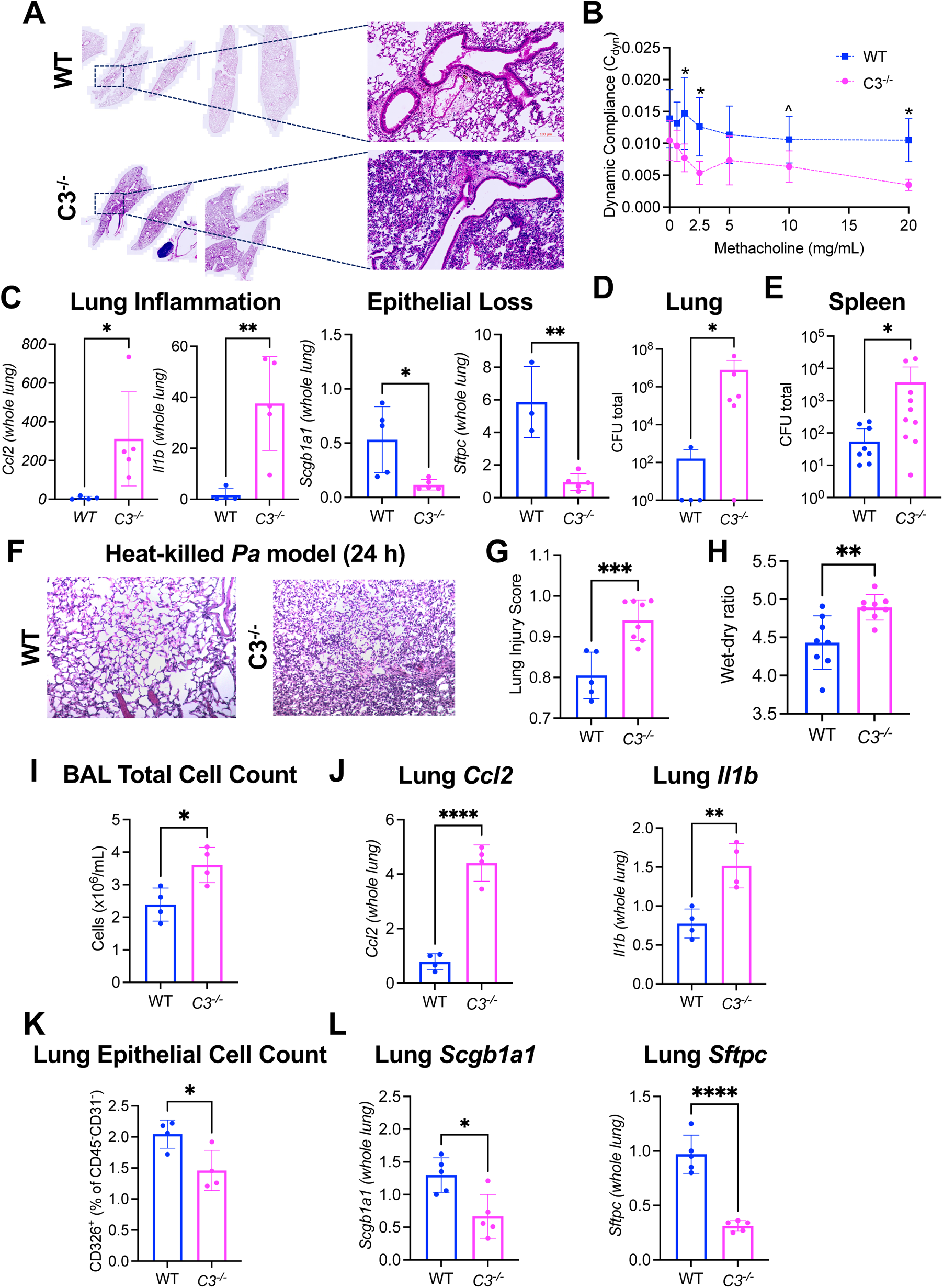
C3 protects against severe bacterial pneumonia. (A-E) Global C3-deficient (*C3^−/−^*) and wildtype (WT) B6 mice were euthanized 24 h after infection with *Pseudomonas aeruginosa (Pa)*. (A) Representative images from n=5 mice/group at 4x magnification; an area of 20x magnification shows both the airways and alveolar space. (B) To assess physiological dysfunction, small animal whole body plethysmography was performed using increasing concentrations of methacholine from 0.625-20 mg/ml (x-axis). Dynamic compliance (in ml/cm H_2_O) represented on the Y-axis. (C) Quantitative reverse transcription polymerase chain reaction (qRT-PCR) measurements for *Ccl2, Il1b, Scgb1a1* (an airway club cell marker) and *Sftpc* (an alveolar type II cell marker), all normalized to *Gapdh*. (D-E) Colony forming units (CFU) in the harvested (D) lungs and (E) spleens. X-axis is on a logarithmic scale. (*F-L*) *C3^−/−^* and WT mice were euthanized 24 h after infection with heat-killed *Pa* (HK*Pa*). (F) Representative histology images at 20x magnification (n=5-8/group). Differences in (G), lung injury scores, (H) lung wet-dry ratios, (I) bronchoalveolar lavage (BAL) cell count, (J) qRT-PCR for *Ccl2* and *Il1b*, (K) proportion of epithelial cells (gated as CD45^-^CD31^-^CD326^+^) from digested mouse lungs, and (L) qRT-PCR for *Scgb1a1* and *Sftpc.* J and L are normalized to *Hprt1*. Distribution of data represented as scatter plots showing individual data points, and bars representing mean ± standard deviation. ^ p < 0.1, * p < 0.05, ** p < 0.01, *** p < 0.001, **** p < 0.0001 using unpaired two-tailed t-test.

One explanation for increased lung injury in C3-deficient mice is damage induced by actively proliferating bacteria. To this end, heat-killed *Pa* (HK*Pa*) were intratracheally instilled into the lungs of both C3-deficient and WT mice, and their lungs and BAL were harvested 24 h later.

Compared to WT, C3-deficient mice still retained more histologic evidence of lung injury (**Fig 1F and 1G**), characterized by an increase in alveolar (**Fig S1I**) and interstitial neutrophilia (**Fig S1J**), and septal thickening (**Fig S1K**). They also had a worse lung wet-dry ratio (**Fig 1H**) compared to WT mice. Similarly, lung inflammation was increased in the C3-deficient mice compared to WT mice, as they had an increased total BAL cell count (**Fig 1I**), and increased lung *Ccl2* and *Il1b* expression post-HK*Pa* administration. Of note, C3-deficient mice treated with HK*Pa* had a lower proportion of epithelial cells on flow cytometry of the lungs (**Fig 1K**), as well as lower expression of *Scgb1a1* and *Sftpc* compared to WT mice treated with HK*Pa* (**Fig 1L**). These observations suggest increased epithelial cell loss post-infection in the setting of C3-deficiency, independent of the bacterial load.

### C3 in the lungs is sourced locally

C3 increases in the BAL of both mice and humans following lung injury (*23*). To determine if this C3 is functional, we adapted our previously established in vitro complement assay, which considers both the activation (using LPS), and thereby, the function of the complement cascade (*24–26*). Functional complement activity in the serum of WT mice was intact both prior to and following 24 h post-*Pa* infection (**Fig 2A**). In comparison to serum, functional complement activity in the BAL of WT mice was upregulated several thousand-fold at 24 h post-*Pa* infection compared to BAL levels in uninfected mice (**Fig 2B**). These observations would suggest that functional complement activity is rapidly induced in an organ with low baseline activity in the first 24 h post infection. To determine the contribution of circulating C3 to that in the BAL, we administered normal mouse serum (NMS, obtained from WT mice) intraperitoneally to C3-deficient mice after intratracheal instillation of *Pa* (to induce injury) or PBS (control). NMS was given either immediately after the *Pa* infection, and the mice were followed for 24 h prior to euthanasia, or it was given 1 h prior to euthanasia, by the time the injury had already occurred. Splenic bacterial dissemination was lower in mice administered NMS compared to C3-deficient serum (**Fig S2A**). C3 was detected in the BAL at 24 h after NMS administration only in the C3-deficient mice administered NMS who had alveolar-capillary barrier disruption in the setting of *Pa* pneumonia, not in those who had only received intratracheal PBS. (**Fig 2C**). However, C3 was not detected in the BAL at 1 h post NMS administration even if the lungs had been injured by *Pa*, despite being detected in the circulation. These observations suggest that following disruption of the alveolar-capillary barrier, circulating C3 entry into the alveolar space is delayed relative to local C3 secretion.

**Figure 2.**
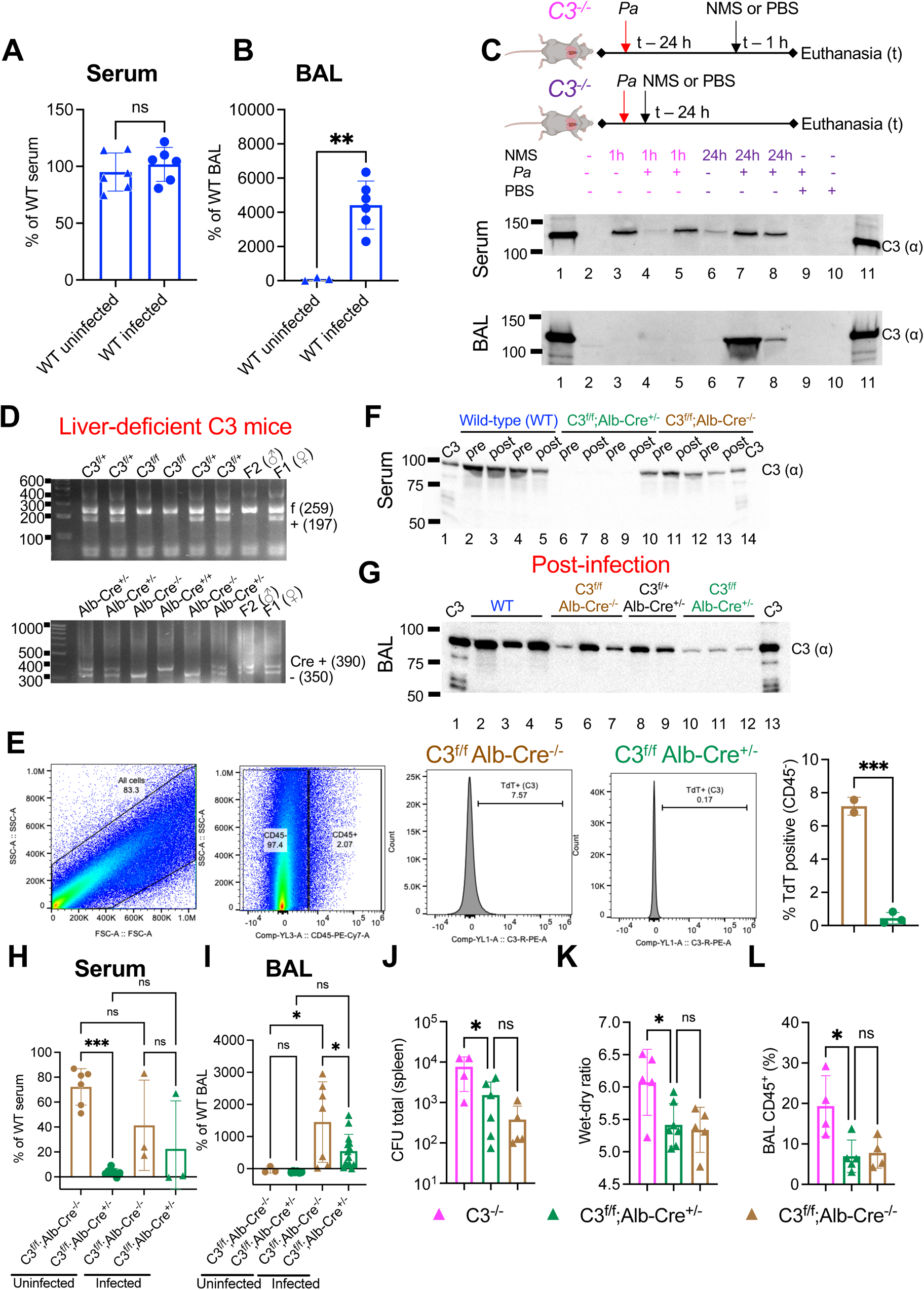
C3 in the lungs is sourced locally. Lipopolysaccharide (LPS)-induced complement activation in (A) mouse serum, and (B) BAL assessed in both uninfected and infected WT mice. Results expressed as % activity of concurrently run uninfected WT B6 mouse sera and BAL respectively. (C) Infected *C3^−/−^* mice were administered normal mouse serum (NMS) or phosphate-buffered saline (PBS) intraperitoneally either 1 h or 24 h before euthanasia (i.e., at the time of infection). (D) PCR products from genomic DNA separated by agarose gel electrophoresis used to identify mice deficient in liver-derived C3 and their littermate controls. (E) Livers from *C3*^f/f^; *Alb*-Cre^+/−^ and *C3*^f/f^; *Alb*-Cre^−/−^ mice analyzed using flow cytometry using negative selection (CD45 excluded immune cells). C3 expression assessed using TdTomato reporter. Immunoblots of reduced (F) serum and (G) BAL from mice drawn pre-and post-*Pa* infection. To assess functional activity in liver-deficient C3 mice, LPS-induced complement activation in (H) mouse serum, and (I) BAL was assessed. Comparison of (J) spleen colony forming units (CFU, x-axis is on a logarithmic scale), (K) lung wet-dry ratios, and (L) BAL immune cell counts post-infection. Distribution of data represented as scatter plots showing individual data points, and bars representing mean ± standard deviation. ns, not significant, * p < 0.05, ** p < 0.01, *** p < 0.001 using unpaired two-tailed t-test.

To quantify C3 expression and production by different cell types, we utilized C3 reporter mice with an IRES (internal ribosome entry side)-tdTomato cassette after the stop signal in the endogenous murine C3 locus, and also has the exons 37-41 sequence flanked by loxP sites enabling its conditional deletion upon the action of a Cre recombinase (*15*). Given that the liver is the major source of circulating C3, we bred these mice to generate an in vivo model of C3 deficiency restricted to the liver (*C3*^f/f^; *Alb*-Cre^+/−^) (**Fig 2D**). We confirmed these mice had no expression of C3 in the liver (**Fig 2E**) and had undetectable levels of circulating C3 compared to littermate controls and WT mice both pre and post *Pa* (**Fig 2F**). However, C3 was detectable (albeit in smaller amounts compared to littermate controls) in the BAL of infected mice at 24 h (**Fig 2G**).

Although there was no functional complement activity in the circulation of the *C3*^f/f^; *Alb*-Cre^+/−^ mice as compared to their littermate controls prior to infection (**Fig 2H**), BAL from *C3*^f/f^; *Alb*-Cre^+/−^ mice was still functional (**Fig 2I**). Post-infection, there was no increase in the functional complement activity in the circulation of the *C3*^f/f^; *Alb*-Cre^+/−^ mice as compared to pre-infection levels (**Fig 2H**). In comparison, the functional activity of the BAL increased at 24 h post-infection in the *C3*^f/f^; *Alb*-Cre^+/−^ mice (**Fig 2I**). Taken together, these observations suggest that although circulating C3 is functional in the lung in the setting of an acute infection, sources of C3 other than the liver also contribute to the C3 function in the lung.

Given the lack of both detectable circulating C3 in the *C3*^f/f^; *Alb*-Cre^+/−^ mice nor any functional complement activity in the serum, we hypothesized that the functional complement activity in the BAL of the *C3*^f/f^; *Alb*-Cre^+/−^ mice was generated from the lung. The presence of a thoroughly functional extra-hepatic source of C3 was corroborated by lower splenic dissemination of bacteria in the *C3*^f/f^; *Alb*-Cre^+/−^ mice as compared to the global C3-deficient mice, but higher in these mice compared to their littermate controls (**Fig 2J**). These findings would suggest that although liver-derived C3 is necessary for limiting bacterial dissemination, sources of C3 other than those from the liver also have an important role. Moreover, there was no difference in either the lung wet-dry ratio (**Fig 2K**), the BAL cell count (**Fig 2L**) or histological evidence of injury (**Fig S2B**) in mice deficient in liver-derived C3 as compared to their littermate controls at 24 h post-*Pa* infection, although it was reduced when compared to global C3-deficient mice. These observations suggest lung-derived C3 played a key role in mitigating bacterial pneumonia-induced lung injury. No differences were observed in BAL cytokine levels in mice deficient in liver-derived C3 as compared to their littermate controls at 24 h post-*Pa* infection (**Fig S2C-F**).

### Non-immune cells contribute to C3 in the lung

To evaluate sources of C3 in the normal human lung, we analyzed publicly available single-cell RNASeq datasets from the Lung Gene Expression Atlas and LungMAP (*27–30*). Our analysis showed that, although most cells expressed C3 transcripts, major sources of C3 in the lungs are epithelial cells, fibroblasts and, to a lesser extent, monocytes/macrophages (**Fig 3A and S3A**).

**Figure 3.**
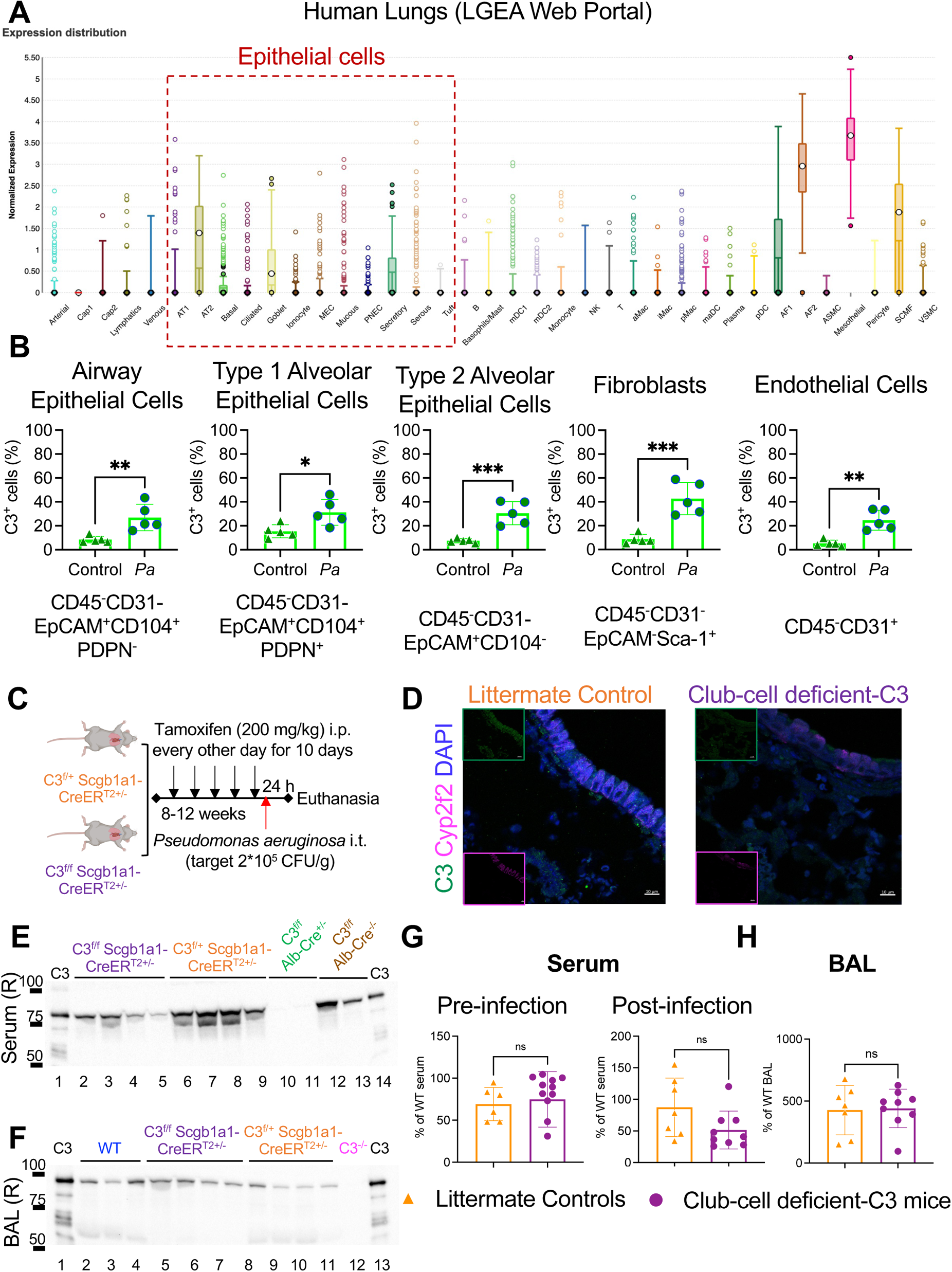
Non-immune cells contribute to C3 in the lung. (A) Human lung expression of C3 (ENSG00000125730) using the LungMAP Single Cell Reference (v1) in the Lung Gene Expression Atlas (LGEA) Web Portal (LungMAP Phase 2). (B) Changes in C3 expression on flow cytometry in cells obtained from lungs of non-infected (Control) and infected (*Pa*) mice deficient in liver-derived C3 (*C3*^f/f^; *Alb*-Cre^+/−^, n=5/group). (C) Schematic for tamoxifen induction of *Cre* recombinase followed by infection. (D) Confocal microscopy imaging of C3 (green, FITC equivalent 488 nm laser) and Cyp2f2 (pink, Cy5 equivalent 640 nm laser) in lungs of *C3*^f/f^; *Scgb1a1*-CreER^T2+/−^ mice, as compared to littermate controls that have normal C3 and Cyp2f2 expression. Cyp2f2 was used as the club cell marker instead of Scgb1a1, as antibodies to both C3 and Scgb1a1 are of the same species (**Table S2**, 63x oil magnification, representative of n=4/group, bar = 10 μ. Individual inserts show single colors. Immunoblots of reduced (E) serum and (F) BAL from lung-deficient C3 mice and littermate controls. Mice deficient in liver-derived C3 and their littermate controls have been used as a comparison for the serum immunoblots, and C3^−/−^ mice have been used to compare BAL. (G) LPS-induced complement activation in mouse serum. (H) The same assay performed on BAL post-infection. Distribution of data represented as scatter plots showing individual data points, and bars representing mean ± standard deviation. ns, not significant, * p < 0.05, ** p < 0.01, *** p < 0.001 using unpaired two-tailed t-test.

To identify cell-intrinsic production of C3 in the lungs in the absence of circulating C3, we profiled C3 producing non-immune cells in liver-deficient C3 mice (*C3*^f/f^; *Alb*-Cre^+/−^) infected with *Pa,* who have an IRES-Td tomato cassette after the stop signal in the endogenous murine C3 locus (C3 expression reporter mice). In these mice, C3 increased by approximately 3-fold in airway epithelial cells and Type 1 alveolar epithelial cells, and increased by nearly 4-fold from baseline in Type 2 alveolar epithelial cells, fibroblasts and endothelial cells at 24 h post-*Pa* infection (**Fig 3B and S3B**). We had previously shown that lung epithelial cells expressing SCGB1A1 (i.e., CCSP) are a major source of C3 in humans, especially in diseases associated with recurrent infections such as cystic fibrosis (*16*), and we were interested in how C3 from cells at the mucosal barrier surface affect the host response to pneumonia. Hence, we generated *C3*^f/f^; *Scgb1a1*-CreER^T2+/−^ mice to specifically delete C3 produced by these epithelial cells (club cell deficient-C3 mice); hereafter, termed as mice deficient in ‘lung-derived C3’ (**Fig 3C and S3C**). Successful deletion of C3 from club cells was confirmed using immunofluorescence for C3 and Cyp2f2, a club cell marker (**Fig 3D**) (*31*). These mice deficient in lung-derived C3 had similar levels of C3 in their circulation and BAL as their littermate controls (**Fig 3E and 3F**). They also had similar serum and BAL functional complement activity compared to their littermate controls (**Fig 3G and 3H**).

### Epithelial (club) cell-derived C3 protects against severe bacterial pneumonia

*C3*^f/f^; *Scgb1a1*-CreER^T2+/−^ mice had worse acute lung injury after *Pa* infection compared to their littermate controls, as represented by measures of alveolar-capillary barrier disruption (**Fig 4A and 4B**). They also had worse histology (**Fig 4C**) and an exaggerated inflammatory response as compared to their littermate controls, as measured by BAL cytokine levels of IL-1β and TNF-α (**Fig 4D-G**). Despite there being no significant differences in the systemic bacterial dissemination (**Fig 4H**), there was increased epithelial cell injury in these mice deficient in lung-derived C3 compared to their littermate controls, as measured by a marked increase in BAL RAGE levels (**Fig 4I**), as well as increased epithelial cell death, as measured by an increase in BAL cytokeratin-18 (CK-18) levels (**Fig 4J**).(*32*) These findings suggest that lung-derived C3 reduces *Pa*-induced lung injury despite adequate circulating (i.e., liver-derived) C3 levels.

**Figure 4.**
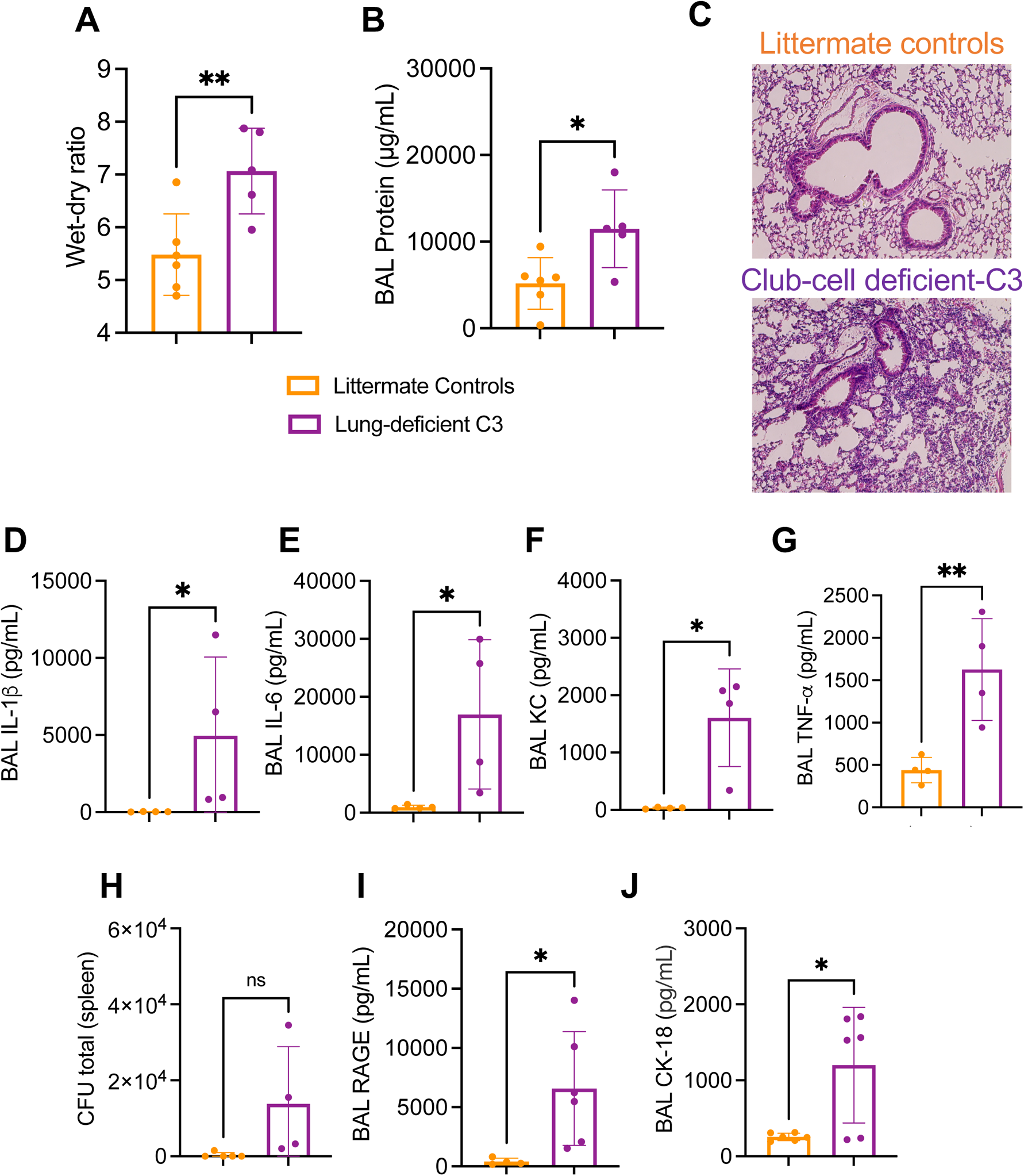
Epithelial cell-derived C3 protects against severe bacterial pneumonia. Mice deficient in epithelial club cell-derived C3 (lung-derived C3, *C3*^f/f^; *Scgb1a1*-CreER^T2+/−^ mice) had worse alveolar-capillary barrier dysfunction at 24 h post-*Pa* infection compared to their littermate controls, as represented by (A) lung wet-dry ratio, and (B) BAL protein. (C) Representative lung histology images of mice deficient in lung-derived C3 and their littermates at 24 h post-infection (20x magnification, n=5/group). (D-G) Comparison of lung inflammation in mice deficient in lung-derived C3 versus their littermates, using cytokines such as interleukin-β), interleukin-6 (IL-6), keratinocyte chemoattractant (KC, CXCL1) and tumor necrosis factor (TNF-α) in the BAL. (H) Comparison of colony forming units (CFU) in the spleens obtained from mice deficient in lung-derived C3 and their littermates at 24 h post-infection. (I) Comparison of epithelial cell injury using the levels of receptor for advanced glycation end products (RAGE) in the BAL of mice deficient in lung-derived C3 and their littermates. (J) Comparison of epithelial cell death using the levels of cytokeratin-18 (CK-18) in the BAL. Distribution of data represented as scatter plots showing individual data points, and bars representing mean ± standard deviation. ns, not significant, * p < 0.05, ** p < 0.01 using unpaired two-tailed t-test.

### Complement Factor B (FB) protects against stress-induced epithelial injury both in vitro and in vivo

Upon activation, C3 is cleaved to form C3a, which binds to the C3a receptor (C3aR), and C3b fragments, which can opsonize targets but also bind to Factor B (FB) to form a C3 convertase (C3bBb) and amplify the alternative pathway of the complement cascade (*33*). To assess which of these putative targets may be facilitating the protective effects of C3 in the lung, we infected global C3aR-deficient (*C3aR^−/−^*) and FB-deficient (*Cfb^−/−^)* mice. C3aR-deficient mice were relatively protected from *Pa*-induced ALI and had a pattern of injury that was similar to the WT mice (**Fig 5A and 1A**). In comparison, FB-deficient mice had more severe injury (**Fig 5A**).

**Figure 5.**
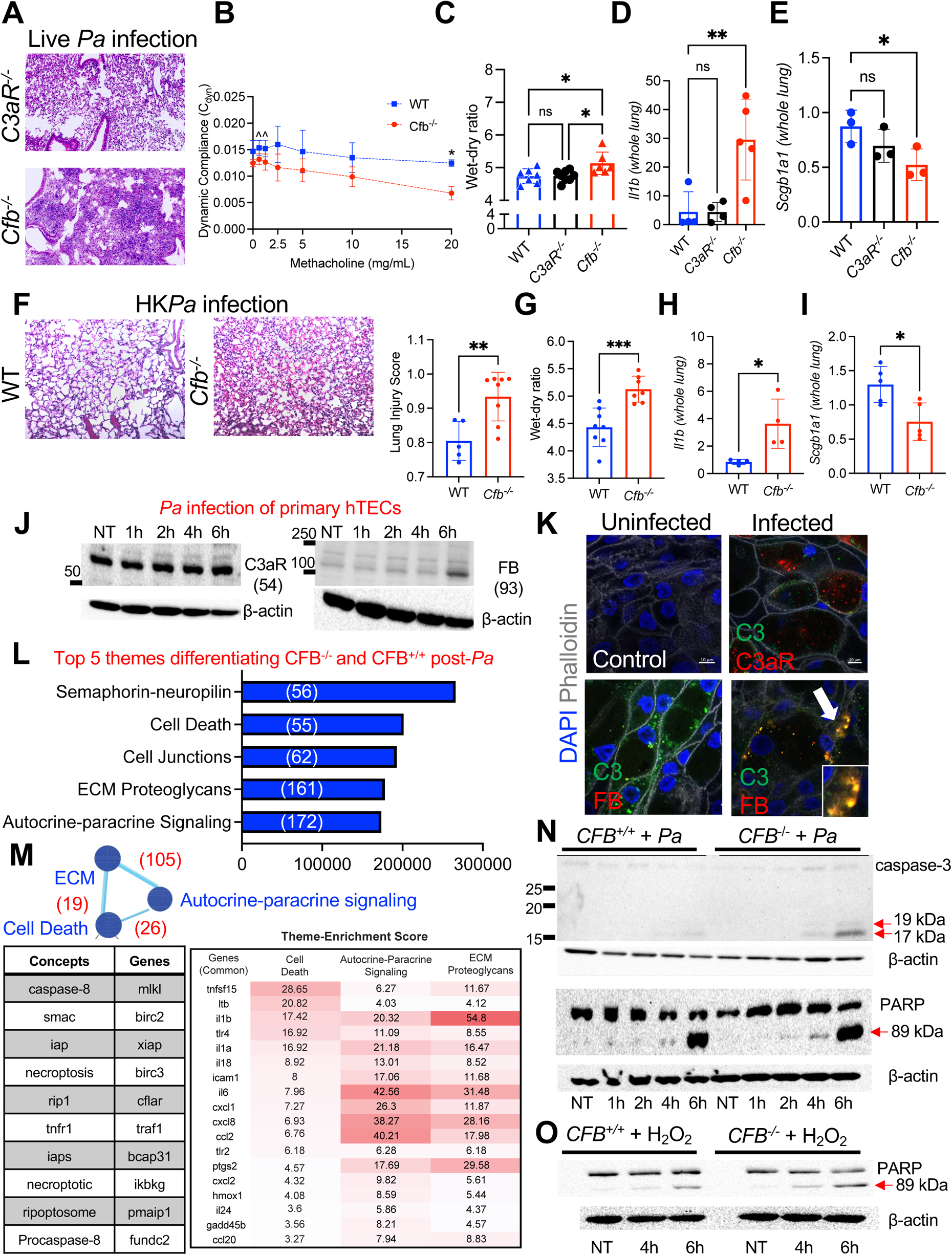
Complement Factor B (FB) protects against stress-induced epithelial injury both in vitro and in vivo. (A-D) Global Factor B-deficient (*Cfb*^−/−^) and C3a receptor-deficient (*C3aR*^−/−^) B6 mice euthanized 24 h after *Pa* infection. (A) Representative images (n=5/group, 20x). These images should be compared to Fig 1A as the lungs were obtained at the same time. (B) Dynamic compliance (y axis, methacholine dose, x axis). (C) Lung wet-dry ratio. (D) *Il1b* expression, normalized to *Hprt1*. (E) *Scgb1a1* expression normalized to *Gapdh.* (F-I) *Cfb*^−/−^ and WT mice administered HK*Pa* and euthanized 24 h after infection. (F) Representative images (n=5/group, 20x) with differences in lung injury scores. These images should be compared to Fig 1F as the lungs were obtained at the same time. Comparison of (G) lung wet-dry ratios, and qRT-PCR for (H) *Il1b*, and (I) *Scgb1a1,* both normalized to *Hprt1*. (J) Immunoblots of FB and C3aR expression in primary human tracheobronchial epithelial cells (hTECs) infected with *Pa* (MOI 5). Representative of n=3 donors. Molecular weights in brackets. (K) Immunostaining for C3, C3aR, and FB in uninfected and infected hTECs using confocal microscopy at 63x (oil) magnification. Phalloidin used to visualize F-actin and identify individual cells. Representative of n=3 donors. (L) A bargraph with the top 5 significant themes plotted by their enrichment scores on RNASeq analysis using the COMPBIO, when comparing infected human BEAS-2B cells (*Pa*, MOI 5, harvested at 4 h) rendered deficient in FB using CRISPR and their FB-sufficient parent clones (the number of genes mapping to these themes are indicated in parentheses. All 5 have p <0.001, which indicates the chance of seeing the theme (genes and concepts) in a random set of genes of the input size). (M) Interrelationship between the three central themes, namely Cell death, ECM proteoglycans and Autocrine-paracrine Signaling. Shared genes between themes are indicated in parenthesis. The top 10 central concepts and genes for the cell death theme are shown in the accompanying table. The pseudo heatmap shows the 18 shared genes common to all three themes and their contribution in each theme. The numeric values indicate the enrichment score of the genes. (N) Immunoblot analysis of caspase-3, PARP and β-actin in FB-deficient human cells and their parent clones (*CFB*^+/+^) after *Pa* infection. Arrows denote cleaved fragments. (O) Similar immunoblot of cleaved PARP in FB-deficient cells and parent clones treated with H_2_O_2_ (250 μM). All qRT-PCR data normalized to *Hprt1.* Distribution of data represented as scatter plots showing individual data points, and bars representing mean ± standard deviation. ^ p < 0.1, * p < 0.05, ** p < 0.01, *** p < 0.001 using unpaired two-tailed t-test for two group comparisons, and ordinary one-way analysis of variance (ANOVA) with multiple comparisons testing for more than two groups.

Increased lung stiffness was observed at 24 h post-infection in FB-deficient mice compared to WT mice, as demonstrated by a worse dynamic compliance (**Fig 5B**) and an increased resistive index (**Fig S4A**). There was increased alveolar-capillary barrier disruption post-infection as evidenced by an increased lung wet-dry ratio in FB-deficient mice compared to WT and C3aR-deficient mice (**Fig 5C**). FB-deficient mice had increased inflammation post-infection, as measured by *Il1b* (**Fig 5D**) and *Ccl2* (**Fig S4B**) expression, and increased number of Ly6C^hi^ monocytes in the lungs of FB-deficient mice compared to WT mice (**Fig S4C)**. FB-deficient mice also had increased epithelial cell loss, measured by lung *Scgb1a1* (**Fig 5E**), *Sftpc* (**Fig S4D**) and *Cdh1* (**Fig S4E)** expression, and increased epithelial cell death compared to WT mice as measured by BAL CK-18 levels (**Fig S4F**) and confirmed using amine dye exclusion on flow cytometry (**Fig S4G**).

A similar pattern of increased lung injury in the setting of FB deficiency was observed in the setting of HK*Pa* administration. FB-deficient mice had worse lung injury on histology (**Fig 5F**), as well as an increased wet-dry ratio as compared to WT mice (**Fig 5G**). These mice also had significantly higher expression of *Il1b* (**Fig 5H**) and *Ccl2* (**Fig S4H**) in their lungs, which paralleled our findings in C3-deficiency. Importantly, FB-deficient mice had decreased *Scgb1a1* (**Fig 5I**), *Sftpc* (**Fig S4I**) and *Cdh1* (**Fig S4J**) expression in their lungs after HK*Pa* administration as compared to WT mice, suggesting epithelial cell loss in the setting of injury.

Given that *Pa* induces a bronchopneumonia, we sought to model these interactions between C3, C3aR and FB in primary human airway epithelial cells. We have previously reported that C3 increases in the primary human tracheobronchial epithelial cells (hTECs) with stress (*16*). In hTECs infected with *Pa*, FB biosynthesis increased post-infection but the levels of C3aR remained unchanged (**Fig 5J**). In line with these observations, C3 colocalized with FB in the setting of an acute *Pa* infection; in contrast, its colocalization with the C3aR was not observed (**Fig 5K**). Hence, we subsequently sought to characterize the consequences of FB-deficiency an C3aR-deficiency in vitro in human lung epithelial cells using CRISPR-Cas9 deletion (**Fig S4K and S4L)**. In an in vitro model of oxidative stress, FB-deficiency was associated with increased cell death compared to the parent clones over time (**Fig S4M**). By comparison, C3aR-deficiency was protective against H_2_O_2_-induced cell death, especially within the first 24 h (**Fig S4N**).

Comparing RNA sequencing (RNA-Seq) of these *Pa* infected FB-deficient cells with their infected FB-sufficient parental line identified cell death as a major (central) theme, utilizing COMPBIO, an ontology-free analysis approach. (www.percayAI.com) (**Fig 5L**). ‘Cell death’ shared common biology with two other themes, ‘ autocrine-paracrine signaling’ and ‘extracellular matrix (ECM) proteoglycans’ (**Fig 5M**), all of which were present in the top 5 enriched themes in the analysis between FB-deficient cells and their parent clones. These three themes shared a common signature consisting of 18 genes, and their contributions towards each theme are highlighted in the pseudoheatmap (**Fig 5M**). Correspondingly, FB-deficiency was associated with increased programmed cell death, evident by increased cleaved caspase-3 and cleaved PARP levels post-*Pa* infection (**Fig 5N**) and was paralleled by similar changes in the setting of oxidative stress (**Fig 5O**). These findings suggest that intracellular FB facilitates lung epithelial cell survival under stress by mitigating programmed cell death.

## DISCUSSION

The effects of the complement system have historically been attributed to its presence in the circulation. However, our understanding of its components being synthesized by tissues and extra-hepatic organs has evolved to recognize that cells at mucosal surfaces may have an intrinsic complement system for host defense (*34–38*). Leveraging advances in complement biology, model systems and technologies, we demonstrate that: (1) C3 protects against acute bacterial pneumonia-induced lung injury; (2) C3 can be sourced locally, independent of that derived from the liver and present in the circulation; (3) structural cells of the lung such as epithelial cells contribute to C3 activity locally; (4) epithelial cell-derived C3 protects against severe bacterial pneumonia; and (5) this protection is potentially occurring via an intracellular complement Factor B (FB), akin to canonical complement activation that occurs extracellularly (*33*), but with important physiological functions necessary for cell survival. These observations put into perspective a long-standing premise that complement — an evolutionary conserved defense system — may have originated intracellularly and evolved extracellularly with the complexity of the host (*39*).

Liver-derived complement proteins form a necessary component of host defense, as evidenced by recurrent infections in patients with cirrhosis (*9, 40*). To that effect, we found that splenic dissemination of bacteria was significantly higher in the mice deficient in liver-derived C3 compared to controls, suggesting its role in enhancing immune resistance. However, there are two specific issues in this context that need consideration: First, that functional complement activity was observed in the bronchoalveolar lavage fluid of mice deficient in liver-derived C3 in the setting of infection, despite no activity in the circulation and, second, that splenic bacterial burden in the mice deficient in liver-derived C3 was significantly lower than that observed in global C3-deficient mice. These observations would suggest that extra-hepatic sources of C3 are induced and are important in controlling bacteremic pneumonia.

We and others have previously reported that lung epithelial cells produce and secrete C3, and this secretion is apically polarized, which may facilitate host defense at the air-liquid interface (*16, 19, 41, 42*). Given our finding that C3 is upregulated in lung epithelial cells under stress in vitro, in the infected airways of people with cystic fibrosis, (*16*), we genetically depleted C3 specifically from lung epithelial cells that are known to be major sources of C3, Scgb1a1^+^ club cells (*43*). The severity of bacterial pneumonia in these mice suggests that lung epithelial cell-derived C3 is necessary for protection against lung injury despite adequate circulating C3 levels. Cell-intrinsic C3 thus affects fundamental cellular processes such as survival, which until now had primarily been demonstrated in vitro (*16, 17*).

C3 is cleaved to C3a and C3b upon cleavage by enzymes. C3a binds to C3aR, functions as an anaphylatoxin and also influences cellular functions such as cytokine production by T cells (*44*). C3b, on the other hand, is as an opsonin but also forms an enzyme called the “C3 convertase” with Factor B (C3bBb) to amplify alternative pathway activation in the circulation (*13, 33*). Our in vitro and in vivo findings indicating that Factor B is necessary for cellular survival, thus, aligns with the concept of an intracellular convertase that has been suggested by others (*19, 45*) and is now strengthened by our findings. Given that C3a-C3aR interactions have previously been shown to facilitate the differentiation of naïve CD4^+^ T cells towards a T_H_1 phenotype (*44*), as well as glucose-induced insulin secretion in human and mouse pancreatic islet cells (*17, 46*), our findings on cytoprotective role of Factor B suggests that intracellular C3 may have multiple targets and functions based on the stressor and the cell type. These observations therefore, create a precedent for complement modulating cellular processes in a stimuli-specific and cell-specific manner, and accordingly targeting therapies based on the desired effect.

There are certain limitations of our study. First, we deleted C3 from Scgb1a1^+^ cells based on our prior findings that (a) these cells are an abundant source of C3 (*16*), (b) C3 forms protects both epithelial and alveolar cells from infection (*47*), and (c) *Pa* infection induces a bronchopneumonia (*48*). Future work will need to address whether the protective effects of lung-derived C3 in severe bacterial pneumonia is due to the biosynthesized C3 that is stored in cells, or is secreted by neighboring lung epithelial cells and is internalized (*16, 21*). Second, there are sources of C3 in the lungs other than lung epithelial cells, such as monocyte/macrophages, fibroblasts and mesenchymal stem cells (*15, 49–51*). Future studies should determine if delivering C3 to the lung via these other sources is sufficient to mitigating lung injury in a paracrine manner. Third, our current in vivo models lack the ability to detect the specific isoform of cytoprotective C3. There is an increasing acknowledgement of a secreted form of C3 that has the signal peptide versus an intracellular cytosolic form with an alternate start site downstream of the signal peptide (*17*). Distinguishing these two distinct forms of C3 may have therapeutic implications if one seeks to to augment C3 within the lung. Additionally, although we have shown that C3 may be exerting its protective effect through FB (a known canonical interaction), we do not exclude non-canonical intracellular interactions, such as how C3 interacts with ATG16L1 in pancreatic β cells (*17*) or in gut epithelial cells, thereby targeting the intracellular bacteria towards autophagolysosomes for degradation (*11*). Finally, we focused on the role of C3 in acute injury due to bacterial pneumonia (i.e., within 24 h). The effects of C3 may be different based on the stressor (*19, 52, 53*) and the temporal course of an infection (*54*).

In summary, we show that in addition to the liver, lung-derived C3 provides host response to pneumonia by promoting both immune resistance and tissue resilience. Among these sources, epithelial-cell derived C3 mitigates severe bacterial pneumonia. This protection appears to occur via Factor B, which is an important component of the alternative pathway of the complement cascade (*55*). Our observation of an intracellular Factor B protein thus adds to the growing knowledge of how intracellular complement proteins modulate fundamental processes such as cell survival (*10, 11, 16, 44, 46*), and importantly, provides a putative mechanism by which lung-derived C3 offers protection. An understanding of tissue-derived and intracellular complement proteins is necessary to reconcile prior trial results of complement therapeutics, a majority of which have primarily targeted circulating complement proteins and do not necessarily penetrate inside the cell (*56, 57*). Thus, we are optimistic that complement therapies targeting a specific organ can be re-evaluated in terms of local protein activities, and caution against the unintentional effects of cell-permeable complement inhibitors. In this manner, we will be able to optimally harness endogenously produced immune mediators including the the complement system to better combat diseases such as pneumonia-induced lung injury.

## MATERIALS AND METHODS

### Materials and Methods

A table following the ARRIVE guidelines for standard reporting of animal studies is included in the Supplementary Material (**Table S1**) (*58*).

### Study Design

#### Research objectives

The primary objective of the research was to assess whether C3-deficiency increases the risk of severe pneumonia occurring in the setting of *Pseudomonas aeruginosa* infection. The secondary objectives included (i) assessing whether mice deficient in liver-derived C3, and mice deficient in lung-derived C3 were at increased risk of severe pneumonia compared to their littermate controls, and (ii) whether the protective effects of C3 occurred via C3aR and/or FB.

#### Research subjects or units of investigation

All animal studies were conducted on protocols approved by the Institutional Animal Care and Use Committee (IACUC) at Washington University School of Medicine (WUSM). Mice (8–12 weeks of age) on a C57BL/6 background were used for all experiments. Commercially available mice (Jackson Laboratories, Bar Harbor, ME) purchased included: wildtype (WT, 000664) and B6.Cg-*Speer6-ps1^Tg(Alb-cre)21Mgn^*/J (Alb-Cre, 003574) mice. *C3*-deficient, *C3aR-*deficient and *Cfb*-deficient mice were maintained in colonies at WUSM (*55, 59, 60*). Scgb1a1-CreER^T2+/−^ mice were generated by breeding an Scgb1a1-CreER^TM^ (B6N.129S6(Cg)-*Scgb1a1^tm1(cre/ERT)Blh^*/J, strain # 016225) to an Ai9 mouse (B6.Cg-*Gt(ROSA)26Sor^tm9(CAG-tdTomato)Hze^*/J, strain# 007909). *C3*^f/f^IRES-tdTomato mice were obtained from Dr. Claudia Kemper at the National Institutes of Health (*15*), and bred with the Alb-Cre mice to create mice deficient in liver-derived C3, and with the Scgb1a1-CreER^T2+/−^ mice to generate mice deficient in lung-derived C3.

Human tissue use has been reviewed by the Institutional Review Board of Washington University. Primary human tracheobronchial epithelial cells (hTEC) and BEAS-2B lung epithelial cells (CRL-9609, ATCC, Mannassas, VA) were used for in vitro experiments. hTECs were isolated from the surgical excess of tracheae and proximal bronchi of lungs donated for transplantation. Tissues from deceased individuals are exempt from regulation by HHS regulation 45 CFR Part 46. The method for cell culture, including media composition, has been described in detail previously (*16*). All cell cultures were maintained at 37 °C in 5% CO2.

C3aR- and FB-deficient BEAS-2B cells were generated by CRISPR nuclease-induced targeted double strand break at the Genome Engineering Center at WUSM. *C3aR* only has one coding exon; hence, the gene was knocked out by using two gRNAs to delete the whole open reading frame (**Fig S4G**). *CFB* was disrupted by introducing out-of-frame indels in all alleles so that the mRNA would be degraded by nonsense mediated decay (**Fig S4H**). These cells were confirmed to be mycoplasma-negative prior to experimentation and were compared to their parent clones in all experiments.

#### Experimental design

All data are reported from controlled laboratory experiments. Mice were infected with *Pseudomonas aeruginosa (Pa)* M57-15, a clinically relevant strain obtained from the sputum of a patient with cystic fibrosis (*61*). The mice were infected with *Pa* for 24 h before euthanasia for blood, bronchoalveolar lavage fluid and tissue harvest. Lungs were harvested either for the wet-dry ratio, BAL collection, flow cytometry, bacterial colony counts, or as a whole organ for fixation in 10% formalin before submission for histology. Spleens were harvested for colony counts. BAL was collected from the mouse lungs using 0.5-1 ml PBS with 1X Halt Protease Inhibitor Cocktail (diluted from 100X; #78429, Thermo Fisher). After collection, the BAL was spun down at 500 g for 5 min before the supernatant was collected and stored at −80 °C for further experiments.

For serum reconstitution experiments, normal mouse serum (NMS, Complement Technology, Inc.) was administered to C3-deficient mice at a dose of 10 μL/g body weight intraperitoneally at the time points noted in the figure legend.

For in vitro experiments, the same *Pa* was used at different multiplicity of infection (MOI) and for different periods of time as indicated in the figure legends. Hydrogen peroxide (H_2_O_2_, H1009, Sigma, St. Louis, MO) was freshly prepared to a concentration of 1000 μM and used for in vitro experiments.

#### Sample size

Mouse experiments included at least 5 mice in each group unless animal death occurred during the procedure. Each experiment was repeated at least twice. The effect was not attributed to the differences between the groups, unless specified. Primary cell culture experiments included at least three different donors. RNA sequencing experiments from BEAS-2B cells included 3 samples in each group.

#### Replicates

Each experiment was performed at least twice, in most cases, by two independent authors.

#### Blinding

Investigators who assessed, measured and quantified the results were blinded to the intervention.

### Experimental Procedures

#### Genotyping

To confirm the genotypes, tips of the tails of the mice were collected before weaning. Oligomers for PCR amplification were designed to yield products of unique size to identify deleted sequences from *C3, C3aR*, and *Cfb* as previously published (*62, 63*). PCR reaction for the digested tail samples is as follows: Step 1 at 94 °C for 4 min, Step 2: 94 °C for 1 min, Step 3: 68 °C for 1 min, Step 4: 72 °C for 1 min, Steps 2-4 repeated for 30 cycles, followed by Step 5 at 72 °C for 5 min, and reaction is concluded at 4 °C until samples are retrieved. Primers used are listed in **Table S3**. The band are visualized with gel electrophoresis using 2-3% agarose (#16500500, Thermo Fisher) gel with SYBR Safe dye (#S33102, Thermo Fisher).

#### Bacterial infection

*Pa* (M57-15 strain) was cultured on plates with LB broth with agar (#L7025-500TAB, Sigma-Aldrich) overnight at 37 °C from frozen stock. Liquid inocula for the mice infections were prepared by inoculating *Pa* from the plates in LB broth (#L3522, Sigma-Aldrich) overnight at 37 °C and keeping in a rotating shaker at 220 rpm for 16 h (logarithmic phase of infection). Subsequently, the bacteria were recovered by centrifugation at 3000 g for 6 min at 4 °C, resuspended in 10 ml sterile PBS, centrifuged again and resuspended a final time in 10 ml PBS. Several dilutions (neat, 1:2, 1:5, 1:10) were prepared in PBS and the absorbance was measured at 380 nm (Epoch Microplate Spectrophotometer, BioTek) to calculate the amount needed to make the required dose using the bacterial growth standard curve.

For in vivo experiments, all mice were infected at a dose of 2 × 10^5^ CFU/g of body weight, with the bacteria being in their logarithmic phase of growth at 16-18 h post-incubation in previously autoclaved Luria broth, and were euthanized at 24 h. To prepare heat-killed *Pa* (HK*Pa*), the stock solution was incubated at 100 °C for 30 min and frozen at −80 °C till use. First, mice were anesthetized with a cocktail of ketamine (100mg/kg) and xylaxine (10mg/kg), intraperitoneally. They were orotracheally intubated utilizing a fiber-optic cable connected to a light source (BioLite Intubation System, BIO-MI-KIT, Braintree Scientific, Braintree, MA to facilitate insertion of a Surflo 20-gauge catheter (# SR-OX2025CA, Terumo Medical Products, Somerset, NJ) within the trachea. A 2×10^5^ CFU/μL suspension of bacteria (or equivalent dose of HK*Pa*) in sterile PBS was introduced into the catheter at a volume of 1 uL per gram of body weight. After 24 h, euthanasia was carried out using avertin (2% working solution of avertin is prepared from 100% avertin, which is made by mixing 1 g of 2,2,2-tribromoethyl alcohol (#T48402, Sigma-Aldrich) with 1 ml of tert-amyl alcohol (#240486, Sigma-Aldrich)),

For in vitro infections, the bacterial cultures were suspended in DMEM (Dulbecco’s Modified Eagle’s Medium, #10-017-CV, Corning, NY) supplemented with 10% heat-inactivated fetal bovine serum (#26140079, Thermo Fisher) containing no antibiotics. *Pa* infection was carried out for different time points at the indicated multiplicities of infection (MOIs) as described in the figure legends.

### Outcome Measures

#### Assessment of functional complement activity in the serum and BAL

We adapted a previously published assay in the serum (*25*). For its use with BAL, ELISA plates (96-well flat-bottom, #3855, Thermo Fisher) were coated with LPS (2 μg/well in 100 μL; #L2762, Sigma-Aldrich) overnight at 4 °C. After washing three times with a solution of 0.05% Tween 20 in PBS, samples (serum 1:10, BAL 1:5) in Mg^2+^-EGTA buffer were added 50μL/well, and incubated at 37 °C for 1 h. The plates were washed again three times and 100μL/well goat anti-mouse C3 (#55463, MP Biomedicals) diluted in 1% bovine serum albumin (BSA, #A7906, Sigma) in PBS (1:4000 in 1%BSA/PBS) was added for 1 h. After another three washes, the samples were incubated with 100μL/well HRP-conjugated donkey anti-goat IgG antibody (1:2000 in 1%BSA/PBS) (#705-035-147, Jackson ImmunoResearch Laboratories) for 1 h. After three washes, TMB Color Substrate (#DY999, R&D Systems) was added 100μL/well and incubated at room temperature (RT) for 10 min. The reaction was stopped by the addition of 50μL/well of 2N H2SO4 (#DY994, R&D Systems), and the OD of the samples were measured at 450 nm (Epoch Microplate Spectrophotometer, BioTek).

#### Assessment of acute lung injury (ALI)

ALI was assessed using multiple domains (*64*). To assess alveolar-capillary barrier disruption, we used the lung wet-dry ratio and/or BAL protein levels. To assess lung inflammation, we used flow cytometry, BAL cytokine level measurements by ELISA, BAL cell counts, or transcriptional expression of certain inflammatory genes (e.g., *Ccl2, Il1β*) measured by qRT-PCR. To assess physiological dysfunction, we performed small animal whole body plethysmography. To assess histological evidence of injury, we obtained images at 20x magnification. To assess for epithelial cell death, we measured death using flow cytometry, BAL RAGE levels using an ELISA, or qRT-PCR expression of certain epithelial cell genes (e.g., *Scgb1a1, Sftpc, Cdh1*).

For the wet-dry ratio, the weight of the “wet” lungs was immediately measured right after euthanasia (#AB54-S, Mettler Toledo, Columbus, OH), and the value obtained was divided by their “dry” weight that was measured after 24 h in an 65 °C oven.

To assess physiological dysfunction, whole body phethysmography was performed. At 24 h post-infection, the mice were anesthetized with the same ketamine/xylazine cocktail as mentioned above and their tracheae were exposed and cannulated with a Surflo 20-gauge catheter (#SR-OX2025CA, Terumo Medical Products). The mice were connected by the cannula to the Buxco FinePointe Resistance and Compliance Station (Data Sciences International, Inc, St. Paul, MN) and their responses to inhaled methacholine (#A2251, Sigma Aldrich) were recorded in a dose-response study, starting from control PBS to increasing doses of nebulized methacholine (0.625, 1.25, 2.5, 5, 10, and 20 mg/ml in PBS), performing deep breaths. Changes in airway resistance (RI) and dynamic compliance (Cdyn) were recorded.

To assess systemic bacterial burden, spleens were harvested into 0.9 ml sterile PBS and homogenized in microtubes (#72.694.006, Sarstedt AG & Co, Nümbrecht, Germany) using stainless steel beads (11079132ss, Biospec, Bartlesville, OK), on a bead beater. The homogenate was serially diluted, plated on LB agar, and incubated at 37 °C overnight prior to colony counting. To assess local bacterial burden, lungs were harvested into 0.8 ml sterile PBS and processed similarly. The total volume of both the spleen and lung homogenate was 1 ml.

#### Flow cytometry

To obtain a single cell suspension mouse lungs were digested using collagenase A (10103586001, Roche, 1.5 mg/ml), DNAse I (D4527, Sigma, 0.1 mg/ml), 5% fetal bovine serum, 10 mM HEPES buffer (25-060-CI, Corning, Durham, NC) and PBS. Specifically, after perfusing the right atrium with 10 ml of 1X PBS, the lungs were inflated with 1 ml of the abovementioned digest solution, cut into 3-4 mm pieces using scissors, and gently vortexed in 2 ml of the digest solution. The digest was incubated at 37 °C for 40 min in a shaker incubator at 190 rpm, while gently vortexing the samples every 5-7 min. After adding 10 ml of ice-cold PBS, the digest was vortexed for 30 seconds, filtered through a 70 µM strainer (431751, Corning) and spun at 500 g for 10 min at 4 °C. After discarding the supernatant, the cells were resuspended in 2 ml ACK Lysing buffer (A10492-01, Gibco) and incubated at RT for 4-7 min, after which they were resuspended in 10 ml of ice-cold 1% BSA in PBS and centrifuged again at 500 g for 10 min at 4° C. 1-2 x 10^6^ cells were added to a 96-well plate for staining using the antibodies listed in **Table S2**. Data was acquired on an Attune Nxt (14 color) laser (Thermo Scientific) and analyzed using FlowJo v10.8.0 (Becton, Dickinson and Company; Ashland, OR).

#### Quantitative reverse-transcriptase polymerase chain reaction

mRNA extraction was performed using the RNAEasy Plus Mini kit (#74134, QIAGEN, Germantown, MD) per the manufacturer’s protocol and cDNA synthesis was performed using the High-Capacity cDNA Reverse Transcription Kit (#4368814, Thermo Scientific) according to manufacturer’s recommendations. cDNA sample was diluted by 5-fold following cDNA synthesis and real-time PCR of each gene was carried out in duplicate on a Roche LightCycler 480 real-time PCR instrument. Each PCR reaction contained 2 µL of cDNA template, 0.6 µL of forward and reverse primers (with the final concentration of 0.5µM/well), 4 µL of the SYBR Green master mix (A46109, Applied Biosystems), and 5.4 µL of RT-PCR grade H_2_O. All primers were ordered from Integrated DNA Technologies (**Table S4**). The PCR program was run as follows: an initial pre-incubation step at 95 °C for 3 min followed by 45 cycles of denaturation at 95 °C for 10 seconds, annealing and elongation at 60 °C for 20 seconds, and the fluorescence reading at 72 °C for 1 second. Ct values were read and exported from the LightCycler 480 software, and the commonly used 2(-ΔΔCt) method was used to calculate relative gene expression using *Gapdh* or *Hprt1* as housekeeping gene(*65*). Data are shown are adjusted to the housekeeping gene.

#### Enzyme-linked immunosorbent assays (ELISA)

For assessing complement activity in the mouse serum and BAL, we adapted the LPS assay as reported in a prior section. Serum was obtained before and after infection from the same mice; however, BAL was obtained from different mice in the uninfected and infected groups, as the BAL procedure is terminal.

For the BAL protein levels (i.e., BCA assay), fresh or thawed BAL samples were diluted (1:10 to 1:50 in PBS) to fall within the range of the recommended standard curve of the assay, before the total protein level was quantified using Pierce™ BCA Protein Assay Kit (#23227, Thermo Fisher) according to provided protocol.

To assess cytokine levels in the mouse serum and BAL, we used the a multianalyte assay (LEGENDplex Mouse Inflammation Panel; #740446, BioLegend, San Diego, CA) according to the provided protocol.

To assess epithelial cell injury in the mouse BAL, the Mouse RAGE DuoSet ELISA (#DY1179, R&D Systems, Minneapolis, MN) was used according to the provided protocol. The BAL samples were either diluted 1:2 or 1:4 based on known protein levels, but in a single run, the dilutions were the same. To assess epithelial cell death in the mouse BAL, the Mouse Cytokeratin 18 ELISA (#ab243678, Abcam, Cambridge, United Kingdom) was used according to the provided protocol. The BAL samples were diluted 1:2 for this assay.

#### Immunoblotting

Samples obtained from mice were diluted in PBS at a ratio of 1:10 for plasma and 2:10 for BAL. Laemmli sample buffer (1610747, Bio-Rad, Hercules, CA) was added to each sample and boiled for 5 min at 100 °C in a heat block for complete denaturation of protein. To prepare cell lysates for immunoblotting, cells were washed in ice-cold PBS and lysed using lysis buffer (9803, Cell Signaling Technology, Danvers, MA) supplemented with Halt Protease Inhibitor Cocktail (78429, Thermo Scientific, Waltham, MA) for 20 min on ice. Lysates were cleared by centrifugation (14,000 g) for 10 min and cooked in SDS-denaturing Laemmli sample buffer for 5 min. Cell lysates were separated on SDS-PAGE (Novex™ WedgeWell™ 4 to 20%, XP04202BOX, Thermo Fisher) in 1X Tris-Glycine-SDS buffer (BP1341-4, Fisher Scientific, Waltham, MA) and transferred using electrophoresis to nitrocellulose membranes by using the iBlot 2 Gel Transfer Device (Thermo Scientific, iBlot 2 NC Regular Stacks, IB23001, Invitrogen, Walham, MA). Membranes were blocked for 1 h at RT in 5% (w/v) nonfat dry milk in TBST (20mM Tris-HCl (pH 7.5), 150mM NaCl and 0.1% Tween 20) and incubated at 4 °C overnight with the antibodies in 5% non-fat dairy milk (w/v) in TBS-T (listed in **Table S2**). For beta actin staining, the membrane was stripped and reprobed. The stripping buffer was prepared by mixing 0.7 ml of β-mercaptoethanol, 2 g SDS and 2.5 ml of 1 M Tris-HCl (pH 6.8) in 100 ml of deionized water. The membrane was incubated for 30 min at 55 °C on a shaker in stripping buffer with gentle agitation followed by washing with TBS-T. The membrane was then blocked and detection with antibody was carried out as described above.

#### Immunostaining

Cells were plated on cover slips in 24 well plates at a seeding density of 10^5^ cells/well overnight prior to the experiment. After treating with *Pa,* cells were fixed by directly adding fixative solution (10% neutral buffered formalin, prepared by mixing 100 ml formaldehyde (37-40%), 900 ml distilled water, sodium phosphate (monobasic) 4 g/L and odium phosphate (dibasic, anhydrous) 6.5 g/L) equal in amount to the medium present in the well. After removing the media and fixative solution after 2 min, 500 µl of fixative solution was again added to the cells and incubation for 20 min at RT. The solution was aspirated and cells were then washed twice with PBS. Cells were permeabilized by adding 0.05% Triton-X 100 (T8787, Sigma, St. Louis, MO) in PBS for 5 min at RT, followed by washing twice with PBS. Cells were blocked by adding 2% bovine serum albumin (9048-46-8, Santa Cruz Biotechnology) in PBS for 30 min at RT followed by washing twice with PBS. After blocking, cells were incubated with primary antibody in 5% donkey serum in 2% BSA in PBS at 4 °C overnight. Cells were subsequently washed thrice with PBS and incubated with secondary antibody at a dilution of 1:200 in 2% BSA for 60 min at RT in the dark. After washing with PBS thrice, the cover slips were mounted using VECTASHIELD Antifade Mounting Medium with DAPI (H-1200, Vector Laboratories, Burlingame, CA). After drying for a few hours, cells were imaged using Zeiss LSM880 laser scanning confocal microscope (Carl Zeiss Inc., Thornwood, NY) at 62X oil magnification.

Immunofluorescence staining of the lungs was performed on paraffin sections. Prior to staining, paraffin embedded lungs tissue sections were deparaffinized using xylene, followed by decreasing concentrations of ethanol. Specifically, slides were dipped in the following sequence for 3 minutes in each solution: 100% xylene, followed by 100% ethanol, 95% ethanol, 75% ethanol, 50% ethanol followed by washing in MilliQ water for hydration and heat-treated with Ag unmasking solution (#H3300, Citrate-Based, Vector Laboratories). Immunostaining was performed with different primary antibodies overnight at 4 °C (**Table S2**). The next day, sections were incubated using Alexa Fluor-conjugated secondary antibodies (**Table S2**) and counterstained using VECTASHIELD Antifade Mounting Medium with DAPI (H-1200, Vector Laboratories, Burlingame, CA). After drying for a few hours, cells were imaged using Zeiss LSM880 laser scanning confocal microscope (Carl Zeiss Inc., Thornwood, NY) at 63X oil magnification.

#### Live-cell imaging

Cell death was analyzed using an IncuCyte Live Cell Imager (Sartorius, Goettingen, Germany) at the Genome Engineering and iPSC Center at Washington University. The ratio of SYTOX Green Nucleic Acid Stain (1:10,000, S7020, Invitrogen) and SYTO 24 Green Fluorescent Nucleic Acid Stain (1:1000, S7559, Invitrogen) was reported over time, averaged over 4 independent fields in each well of a 96-well plate, with 4 technical replicates per time point.

#### RNA extraction and sequencing

RNA was isolated from lung epithelial cells by using the RNeasy plus Mini Kit (74134, QIAGEN). The starting number of cells used for extraction was about 8 × 10^6^. Cells were washed twice with ice-cold PBS, scraped in 1 ml of PBS and centrifuged at 3000 x g for 5 min at 4 °C. The pellet was homogenized using the kit reagents and processed as per the insert. Isolated RNA was sent to Genome Technology Access Center at the McDonnell Genome Institute (GTAC@MGI), WUSM for RNA sequencing. Poly-A selection was used for library preparation and sample processing. A target of at least 30 million reads per library utilizing NovaSeq S4, 2×150 reads was accomplished. Dual indexing during library construction was used to uniquely identify each sample.

#### Analysis of RNA Sequencing data from BEAS-2B cells

Differentially expressed genes (1.6 fold, adjusted p value <0.05) were further analyzed by a novel knowledge engine called ‘COmprehensive Multi-omics Platform for Biological InterpretatiOn’ (COMPBIO V2.0) (*66*). COMPBIO performs a literature analysis to identify relevant processes and pathways represented by differentially expressed, or otherwise related, biological entities (genes, proteins, miRNA’s, or metabolites) or their synonyms using Pubmed abstracts. Knowledge is extracted in an ontology-free manner and maintained in a Persistent Contextual Memory Model (PCMM) that can be aggregated, normalized and computed across from any stored dimension. Conditional probability analysis is utilized to compute the statistical enrichment of contextually associated biological concepts (processes/pathways) over those that occur by random sampling. These related concepts built from the input entities are further clustered into higher-level themes (e.g., biological pathways/processes, cell types and structures, etc.).

Genes, concepts, and theme enrichment is calculated using a multi-component function referred to as the Normalized Enrichment Score (NES) which reflects both the rarity of the concept event associated with an entity list, as well as its degree of overall enrichment. Entity-concept scores above 10.0, 100.0, and 1,000.0 are labeled as moderate, marked, or high in level of enrichment above random. Theme scores are then computed from their constituent entity-concept NES scores. As such, theme enrichment scores above 10,000, 20,000, and 30,000 are identified as having moderated, marked and high levels of entity-concept enrichment, respectively.

Concepts are biologically relevant words or names that have been collected from a multitude of sources of biological language (text) or entities. Within COMPBIO themes, concepts and their relationships are computed in a fully automated fashion. The themes generated by COMPBIO can be annotated with a summary level description of the biological processes and pathways that they represent. This can be achieved either by semi-automated annotation or human assessment. Semi-automated annotation occurs through a comparison of, up to, the 10 most central concepts of each theme with pathways and processes in common knowledge bases such as GO, Reactome, and several others. The most closely associated entries are computed and presented, with score, for annotation of theme. However, as COMPBIO is not restricted to these knowledge bases in its biological discovery process, some themes can only be annotated with human assessment. In these cases, the process is similar in that the annotation is typically a summary description of the biology described by the most central concepts to the theme.

#### Single cell RNA sequencing analysis (scRNASeq) from LungMAP

The single cell reference program for human lungs was used on the Lung Gene Expression Atlas (LGEA, LungMAP Phase 2) web portal (*27–30*), available at: https://research.cchmc.org/pbge/lunggens/tools/lung_at_glance.html?tab=reference.

Cell ontologies are obtained from the NHLBI Molecular Atlas of Lung Development Program Consortium (*67*). The LungMAP scRNASeq reference is associated with the Lung CellCards resource (*30*). This initial reference integrated 259k cells from 72 donors from five published (*68–72*) and one unpublished single cell RNA-seq cohort. It consists of non-diseased adult and pediatric healthy lung single-cell 10x Genomics captures (v2 and v3).

### Statistical Analysis

Data are presented as scatter plots showing individual data points, and dispersion is shown by mean ± standard deviation. The unpaired t test was used for comparing two groups, assuming equal variance and the ordinary one-way ANOVA test with multiple comparisons testing was used for more than two groups. A p value of < 0.05 was considered significant, although values > 0.05 have been reported when there was evidence of a trend. Prism v9.3.0 (GraphPad, San Diego, CA) was used for statistical analysis.

## SUPPLEMENTARY MATERIALS

**Supplementary Figure 1. C3 protects against severe bacterial pneumonia.**

**Supplementary Figure 2. Deficiency in liver-derived C3 did not increase the predisposition to *Pseudomonas aeruginosa (Pa)*-induced acute lung injury.**

**Supplementary Figure 3. C3 in the lungs is produced locally.**

**Supplementary Figure 4. Complement Factor B (FB) protects against stress-induced epithelial injury both in vitro and in vivo.**

**Supplementary Table S1. ARRIVE Guidelines checklist for animal research.**

**Supplementary Table S2. Antibodies used for experiments.**

**Supplementary Table S3. Primers used for genotyping of mice.**

**Supplementary Table S4. Primers used for qRT-PCR of mouse lungs.**

**Table S1.** ARRIVE Guidelines checklist for animal research.

|  | **Item** | **Section/line number, or reason for not reporting** |
| --- | --- | --- |
| 1. | **Study Design** | A general description of all strains used are provided in the section “Research subjects or units of investigation” under Methods. The groups being compared, including control groups, number of animals and the experimental units are reported in the Figure Legends for each figure. |
| 2. | **Sample Size** | A general description of sample size has been provided in the section “Sample size.” under Methods. The number of animals and the experimental units are reported in the Figure Legends for each figure. |
| 3. | **Inclusion and Exclusion Criteria** | For *C3<sup>-/-</sup>* mice, males were used as reported in Figure S1. For *C3aR<sup>-/-</sup>* and *FB<sup>-/-</sup>* mice, there was no effect of sex on the phenotype and hence, animals of both sexes were used. |
| 4. | **Randomisation** | No specific randomization schema was used. However, to minimize potential confounders, mice were infected and euthanized alternately (e.g., Group 1 mouse 1, Group 2 mouse 1, Group 1 mouse 2, Group 2 mouse 2...) |
| 5. | **Blinding** | The person collecting the data (i.e. outcome assessment) for each experiment was blinded to the group allocation. |
| 6. | **Outcome Measures** | Listed under the “Outcome Measures” section in the Methods. |
| 7. | **Statistical methods** | Listed under the “Statistical Analysis” section in the Methods. |
| 8. | **Experimental Animals** | Species-appropriate details have been provided under “Research subjects or units of investigation” in the Methods section. |
| 9. | **Experimental Procedures** | Detailed procedures to facilitate replication, including rationale, have been described under the section “Experimental Procedures” in the Methods section. |
| 10. | **Results** | All figures have been represented as scatter plots with individual data points, and range is represented as mean ± standard deviation. |

**Table S2.** Antibodies used for experiments

| **Antibody** | **Supplier** | **Catalog #** | **Dilution** |
| --- | --- | --- | --- |
| **Immunoblotting** |  |  |  |
| Goat anti-C3 | MP Biomedicals | 55444 | 1:3000 |
| Goat anti-Factor B | Complement Technology, Inc. | A235 | 1:3000 |
| Rabbit anti-caspase 3 | Abcam | ab13847 | 1:1000 |
| Rabbit anti-PARP | Cell Signaling Technology | 9542 | 1:1000 |
| Mouse anti-beta actin | Santa Cruz Biotechnology | sc-47778 | 1:2000 |
| Anti-goat IgG, HRP-linked | Thermo Fisher | 31402 | 1:4000 |
| Anti-mouse IgG, HRP-linked | Cell Signaling Technology | 7076 | 1:4000 |
| Anti-rabbit IgG, HRP-linked | Cell Signaling Technology | 7074 | 1:4000 |
| **Immunofluorescence** |  |  |  |
| Chicken Anti-C3 (human) | Sigma | GW20073 | 1:100 |
| Rabbit anti-C3aR (human) | Invitrogen | PA5-29979 | 1:100 |
| Goat Anti-Factor B (human) | Quidel | A311 | 1:100 |
| Rabbit anti-C3 (mouse) | Antibody Verify | AAS72600 | 1:50 |
| Rabbit anti-Scgb1a | Seven Hills Bioreagents | WRAB-3950 | 1:100 |
| Rabbit anti-Prosurfactant Protein C (proSP-C) | Millipore Sigma | AB3786 | 1:100 |
| Mouse anti-Cyp2f2 | Santa Cruz | sc-374540 | 1:100 |
| Goat anti-Chicken IgY (H+L) Secondary, Alexa Fluor 647 | Thermo Fisher | A-21449 | 1:300 |
| Donkey anti-Goat IgG (H+L) Cross-Adsorbed Secondary, Alexa Fluor 555 | Thermo Fisher | A-21432 | 1:300 |
| Donkey anti-Rabbit IgG (H+L) Cross-Adsorbed Secondary, Alexa Fluor 488 | Thermo Fisher | A-21206 | 1:300 |
| Donkey anti-Rabbit IgG (H+L) Cross-Adsorbed Secondary, Alexa Fluor 555 | Thermo Fisher | A-31572 | 1:300 |
| Goat anti-Mouse IgG (H+L), Cross-Adsorbed Secondary, Alexa Fluor 647 | Thermo Fisher | A-21235 | 1:300 |
| **Flow Cytometry** |  |  |  |
| BV711 anti-mouse CD45 | Biolegend | 103147 | 1:200 |
| APC/Cy7 anti-mouse CD31 | Biolegend | 102440 | 1:100 |
| BV510 anti-mouse CD326 (EpCAM) | Biolegend | 118231 | 1:100 |
| APC anti-mouse CD104 | Biolegend | 123612 | 1:100 |
| PerCP/Cy5.5 anti-mouse PDPN (podoplanin) | Biolegend | 127422 | 1:100 |
| PE-Cy7 anti-mouse Ly-6A/E (Sca-1) | eBioscience | 25-5981-81 | 1:100 |
| Zombie (Zombie Violet Fixable Viability Kit) | Biolegend | 423114 | 1:400 |

**Table S3.**
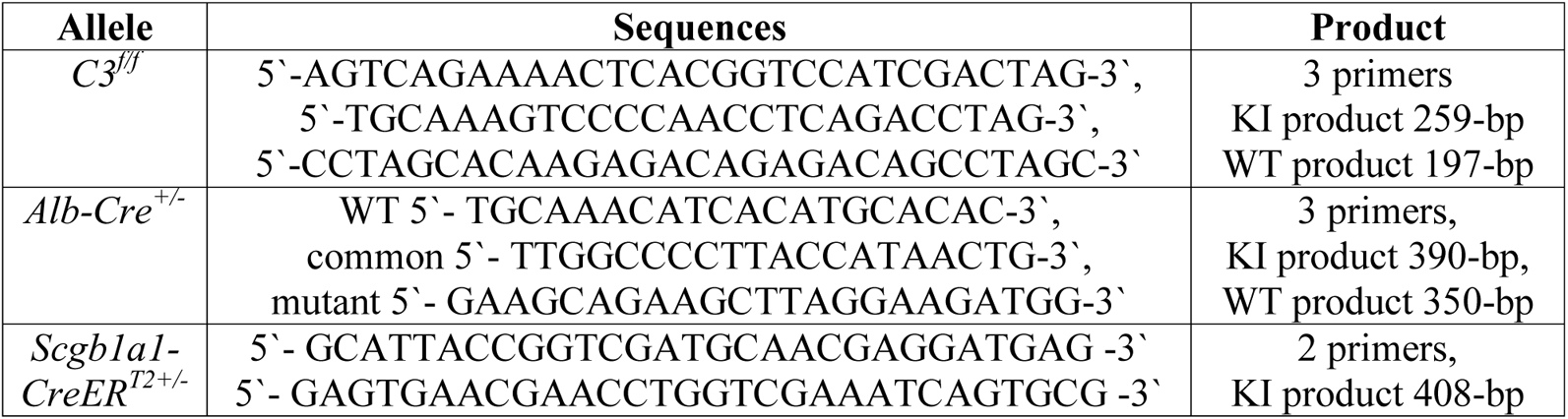
Primers used for genotyping of mice.

**Table S4.**
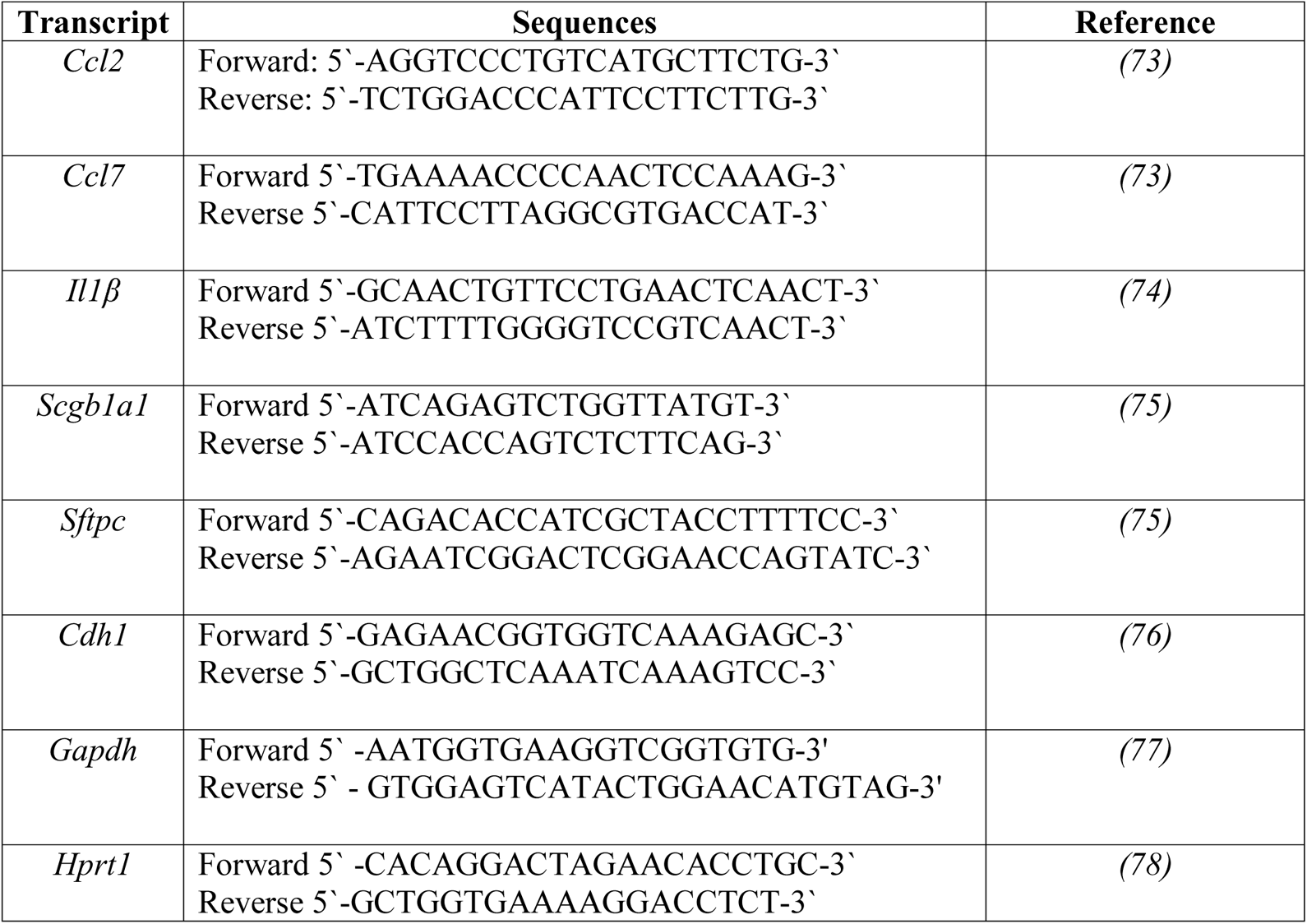
Primers used for qRT-PCR of mouse lungs.

## Acknowledgements

We thank Drs. Claudia Kemper, Dennis Hourcade, Jeffrey Haspel, Michael Holtzman, Jeffrey Koenitzer and Yi-Chieh Perng for their insights on the work and contribution of reagents. We thank Lynne Mitchell, Kathy Liszewski, Carrie Gierasch and Abigail Pesavento for technical assistance.

## Funding

Experimental support was also partially provided by the Bursky Center for Human Immunology and Immunotherapy Programs at Washington University Immunomonitoring Laboratory, in support of the Rheumatic Diseases Core Center (NIH WLC6313040077).

National Institutes of Health grant K08HL148510 (HSK)

National Institutes of Health grant R35GM136352 (JPA)

National Institutes of Health grant R01EYE028602 (JPA)

Children’s Discovery Institute (HSK)

Rheumatic Diseases Research Resource-Based Center (P30AR073752, HSK).

The Clark Family/Clayco Foundation International Cerebroretinal Vasculopathy (CRV) Research Award (JPA)

The content is solely the responsibility of the authors and does not necessarily represent the official view of the NIH.

## Author contributions

Conceptualization: HSK, SLB, JPA

Methodology: ANO, DHK, SKS, LD, LM, JM, LG, JK and XW

Investigation: HSK, ANO, DHK and SKS

Visualization: HSK, ANO, DHK, RAB and SKS

Funding acquisition: HSK and JPA

Project administration: HSK

Supervision: HSK and JPA

Writing – original draft: HSK, ANO, SKS, JPA and SLB

Writing – review & editing: HSK, ANO, DHK, RAB, SKS, LD, LM, JM, LG, JK, XW, JPA, SLB

## Data and materials availability

All data needed to evaluate the reported conclusions are present in the paper or the Supplementary Materials. The RNASeq dataset will be available at NCBI GEO upon peer review publication https://www.ncbi.nlm.nih.gov/geo/).

## SUPPLEMENTARY FIGURE LEGENDS

**Supplementary Figure 1. C3 protects against severe bacterial pneumonia.** (A) To assess physiological dysfunction in C3-deficient mice (*C3^−/−^*) and wildtype (WT) mice, small animal whole body plethysmography was performed using increasing concentrations of methacholine from 0.625-20 mg/ml (x-axis) at 24 h post-*Pa* infection. Airway resistance (invasive resistance, RI, in cmH_2_O.s/ml, y axis). Data points and error bars represent mean ± standard deviation (n=4-6/group). Alveolar-capillary barrier disruption assayed by the lung wet-dry ratio in (B) *C3^−/−^* and WT mice, and (C) C3-deficient male (♂*C3^−/−^*) versus C3-deficient female (♀*C3^−/−^*) mice post-*Pa*. Lung inflammation, determined as (D) the proportion of lung Ly6C^hi^ monocytes (n=4/group), and BAL cytokines such as (E) chemokine ligand 2 (CCL2), (F) interleukin-1β (IL-1β), (G) interleukin-6 (IL-6), and (H) tumor necrosis factor (TNF-α). (I-M) Histology assessed using individual components of a lung injury score. Distribution of data represented as scatter plots showing individual data points, and bars representing mean ± standard deviation. ns, not significant, * p < 0.05, ** p < 0.01 using a unpaired two-tailed t-test, except for (B) and (D-H), a one-tailed t-test was used given the null hypothesis was that C3-deficient mice had worse lung injury than WT mice, based on the initial findings.

**Supplementary Figure 2. Deficiency in liver-derived C3 does not increase the predisposition to *Pseudomonas aeruginosa (Pa)*-induced acute lung injury.** (A) Spleen CFU comparing C3^−/−^ mice receiving C3^−/−^ serum versus C3^−/−^ mice receiving normal mouse serum (NMS or C3-sufficient serum) immediately after *Pa* infection, 24 h prior to euthanasia. (B) Representative lung histology images of mice deficient in liver-derived C3 and their littermates at 24 h post-infection (n=4/group). (C) To assess lung inflammation, we compared the levels of cytokines such as interleukin-1β (IL-1β), interleukin-6 (IL-6), keratinocyte chemoattractant (KC, i.e., CXCL1) and tumor necrosis factor (TNF-α) in the BAL of mice deficient in liver-derived C3 versus their littermates. Distribution of data represented as scatter plots showing individual data points, and bars representing mean ± standard deviation. ns, not significant using a unpaired two-tailed t-test.

**Supplementary Figure 3. C3 in the lungs is produced locally.** (A) C3 (ENSG00000125730) expression in query cells and annotations projected onto the reference UMAP on the LGEA web portal, using the LungMAP Single Cell Reference (v1). (B) Gating strategy for flow-cytometric analysis of cells derived from the lungs of mice deficient in liver-derived C3. (C) Agarose gel electrophoresis of PCR products used to identify genotypes of mice used in experiments. PCR products were imaged using SYBR green and size indicated by notation on the right. Gel showing PCR products from tails of *C3*^f/f^; *Scgb1a1*-CreER^T2+/−^ mice (i.e., mice deficient in lung-derived C3), and their littermate controls (*C3*^f/+^; *Scgb1a1*-CreER^T2+/−^ and *C3*^f/f^; *Scgb1a1*-CreER^T2-/-^ mice). Abbreviations: AF1: alveolar fibroblast 1; AF2: alveolar fibroblast 2; aMac: alveolar macrophage; ASMC: airway smooth muscle cell; AT1: Type 1 alveolar epithelial cells, AT2: Type 2 alveolar epithelial cells; Cap1: capillary cell 1; Cap2: capillary cell 2; abbreviations for two capillary subsets; DC: dendritic cells; iMac: interstitial macrophages; mDC1: myeloid type 1 dendritic cell; maDC: mature human dendritic cell; mDC2: myeloid type 2 dendritic cell; MEC: myoepithelial cells; NK: NK cells; pDC: plasmacytoid dendritic cells; PDPN: podoplanin; pMac: PNEC: pulmonary neuroendocrine cell; SCMF: Secondary crest myofibroblast cell; VSMC: vascular smooth muscle cell.

**Supplementary Figure 4. Complement Factor B (FB) protects against stress-induced epithelial injury in vitro and in vivo.** (A) Airway resistance (y-axis) measured using whole body plethysmography post *Pa*-infection. Data points and error bars represent mean ± standard deviation (n=3-5/group). Lung inflammation post-infection assessed using (B) qRT-PCR of *Ccl2,* and (A) proportion of Ly6C^hi^ monocytes using mass cytometry. Epithelial cell loss assessed using qRT-PCR of (D) *Sftpc,* and (E) *Cdh1* (cadherin), and death assessed by (E) BAL CK-18, and (F) epithelial cells (CD45^-^CD31^-^CD326^+^Zombie^+^) on flow cytometry. (H) Lung *Ccl2* expression to assess inflammation post-HK*Pa*. Lung (I) *Sftpc* and (J) *Cdh1* expression to assess epithelial cell loss post-HK*Pa*. Schematic for CRISPR-induced CRISPR-induced (K) FB (*CFB)* deletion, and (L) C3aR (*C3aR1)* deletion in BEAS-2B cells. Quantification of SYTOX^+^:SYTO24^+^ cells on live cell imaging (Incucyte) over time when (M) *CFB^−/−^* cells and their parent clones (*CFB^+/+^*), and (N) *C3aR^−/−^* cells and their parent clones (*C3aR^+/+^*) were treated with hydrogen peroxide (H_2_O_2_). B and D normalized to *Gapdh.* C, I and J normalized to *Hprt1*. Distribution of data represented as scatter plots showing individual data points, and bars representing mean ± standard deviation. * p < 0.05, ** p < 0.01, *** p < 0.001 using unpaired two-tailed t-test for two group comparisons, and ordinary one-way analysis of variance (ANOVA) with multiple comparisons testing for more than two groups.

