## SUPPLEMENTARY FIGURE LEGENDS for "Lung epithelial cell-derived C3 protects against pneumonia-induced lung injury"

### Figure S1

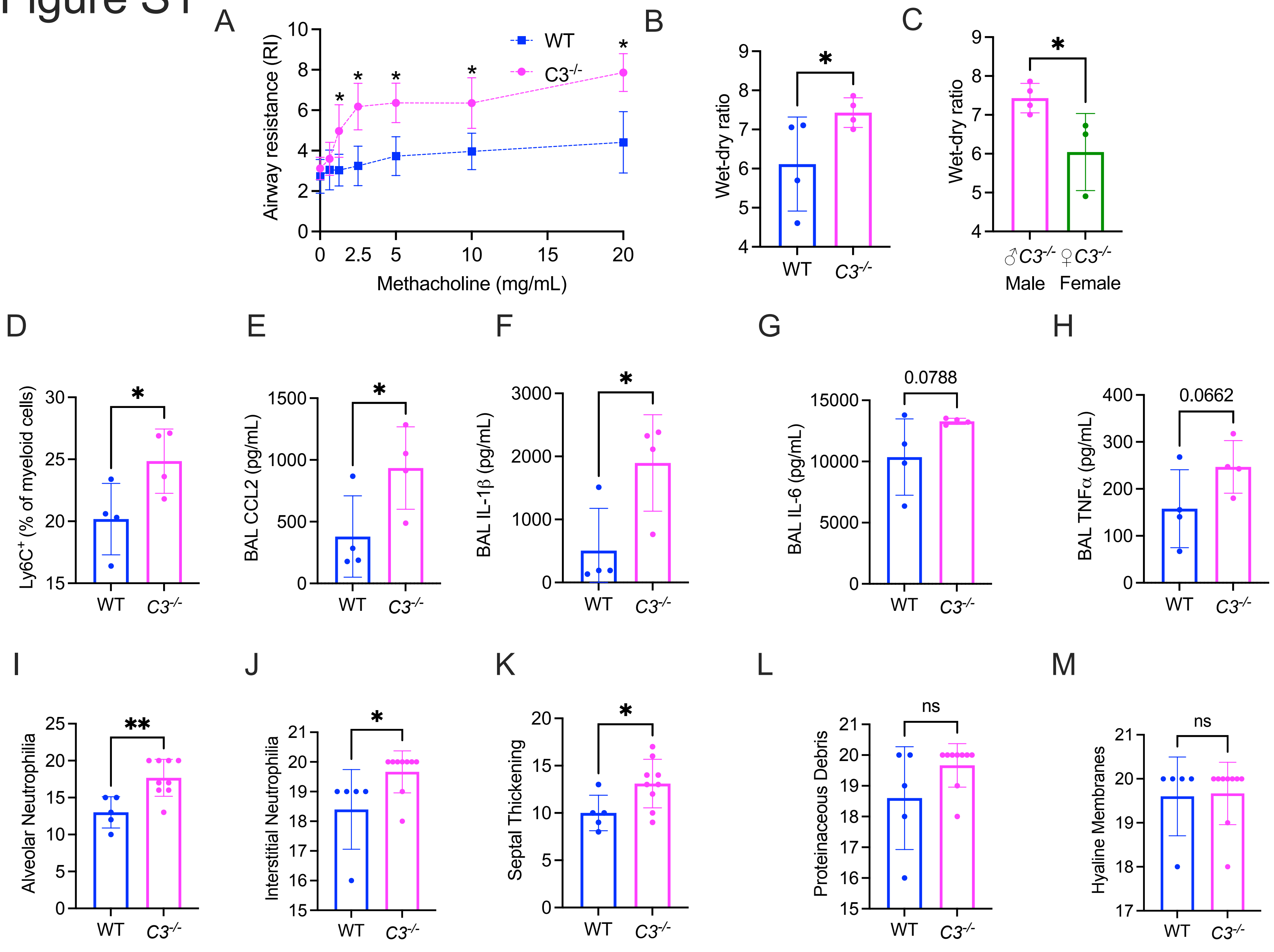

A
Airway resistance (RI)
Methacholine (mg/mL)
WT
C3-/-
B
Wet-dry ratio
WT
C3-/-
C
Wet-dry ratio
♂C3-/- Male
♀C3-/- Female
D
Ly6C+ (% of myeloid cells)
E
BAL CCL2 (pg/mL)
F
BAL IL-1β (pg/mL)
G
BAL IL-6 (pg/mL)
0.0788
H
BAL TNFα (pg/mL)
0.0662
I
Alveolar Neutrophilia
J
Interstitial Neutrophilia
K
Septal Thickening
L
Proteinaceous Debris
ns
M
Hyaline Membranes
ns

### Figure S2

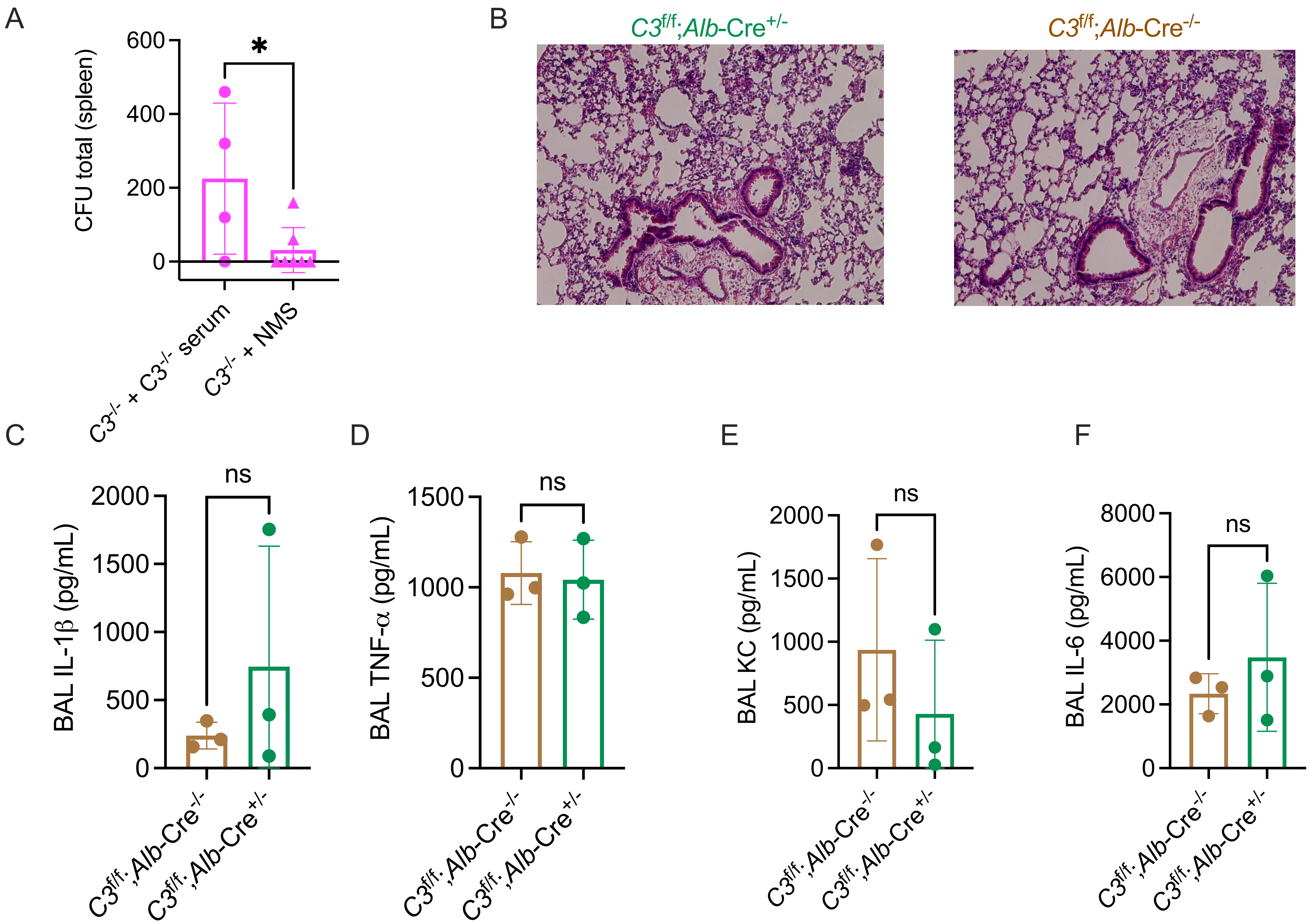

A
600
400
200
0
CFU total (spleen)
*
C3-/- + C3-/- serum
C3-/- + NMS
B
C3f/f;Alb-Cre+/-
C3f/f;Alb-Cre-/-
C
ns
2000
1500
1000
500
0
BAL IL-1β (pg/mL)
C3f/f;Alb-Cre-/-
C3f/f;Alb-Cre+/-
D
ns
1500
1000
500
0
BAL TNF-α (pg/mL)
C3f/f;Alb-Cre-/-
C3f/f;Alb-Cre+/-
E
ns
2000
1500
1000
500
0
BAL KC (pg/mL)
C3f/f;Alb-Cre-/-
C3f/f;Alb-Cre+/-
F
ns
8000
6000
4000
2000
0
BAL IL-6 (pg/mL)
C3f/f;Alb-Cre-/-
C3f/f;Alb-Cre+/-

### Figure S3

A

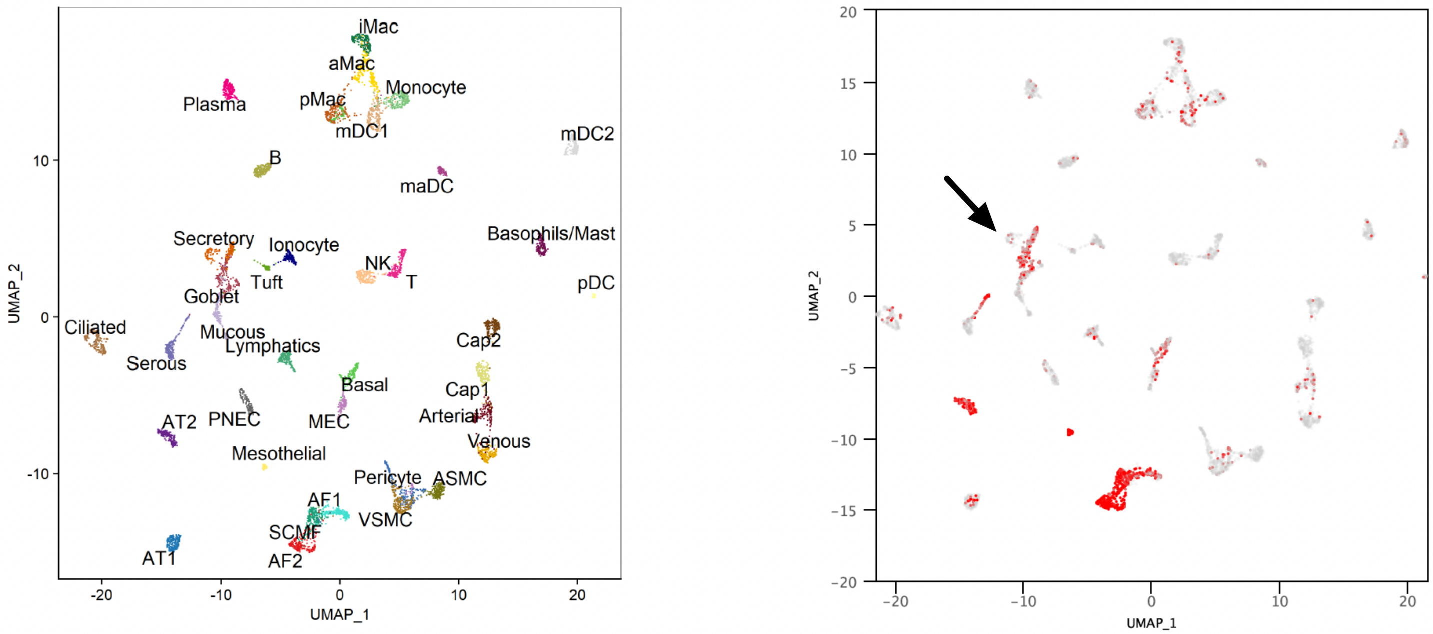


B

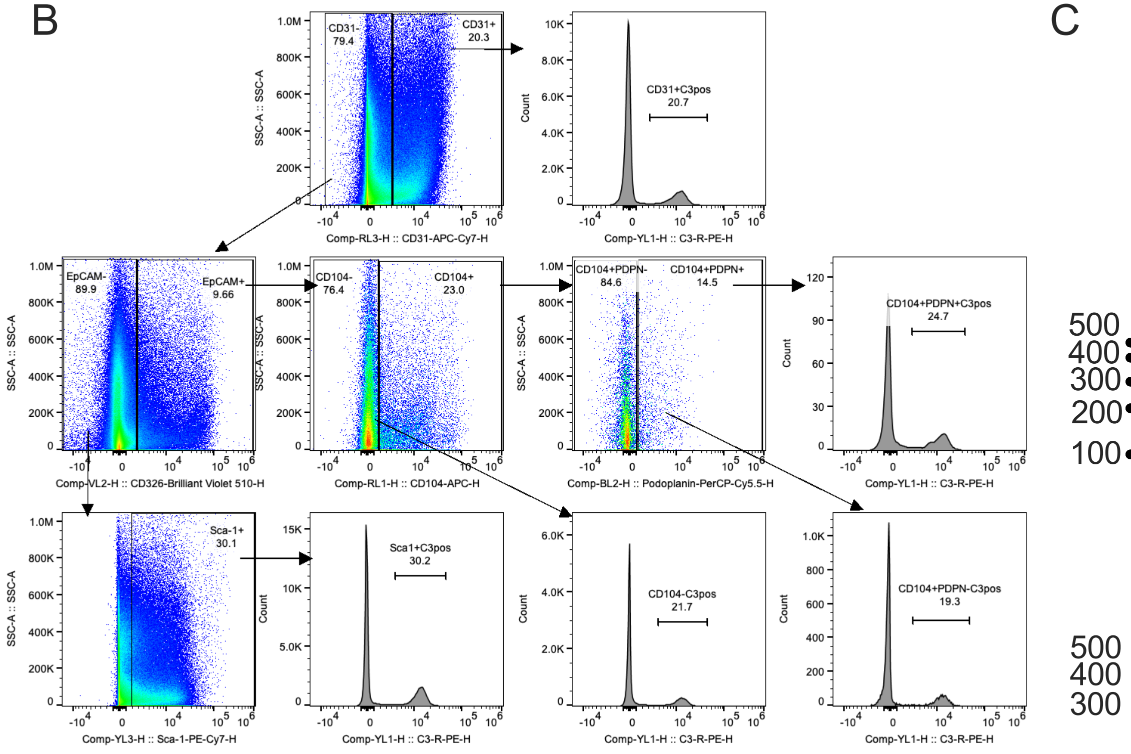


Gating Strategy for Lungs Obtained from *C3*$^{f/f}$;*Alb*-Cre$^{+/-}$ Mice at 24 h Post-Infection (CD45$^{-}$ population)

C

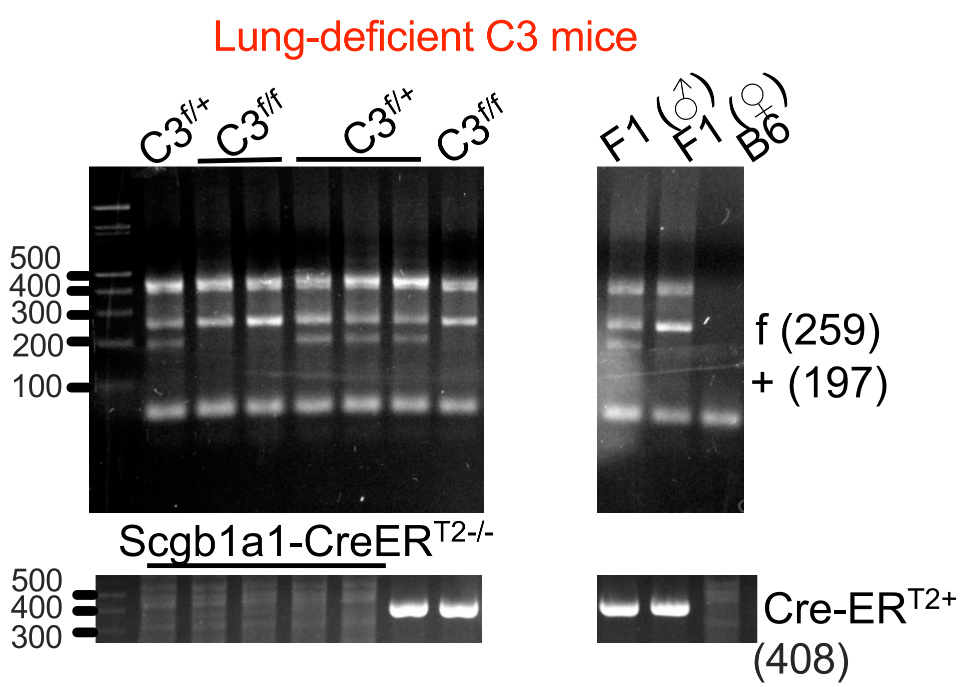

### Figure S4

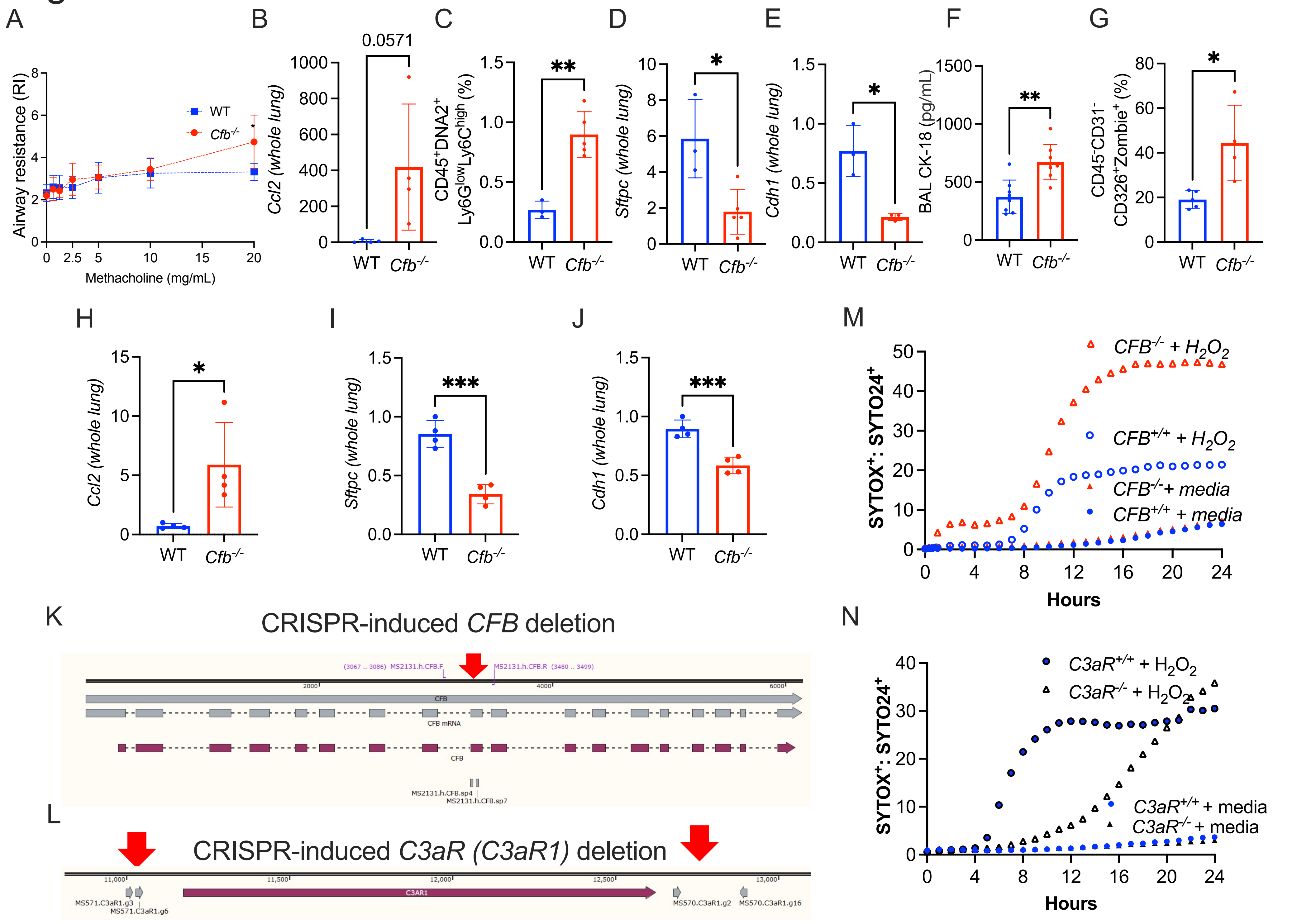

A
B
C
D
E
F
G
Airway resistance (RI)
Methacholine (mg/mL)
WT
Cfb-/-
0.0571
Ccl2 (whole lung)
WT
Cfb-/-
CD45+DNA2+ Ly6GlowLy6Chigh (%)
**
Sftpc (whole lung)
*
Cdh1 (whole lung)
*
BAL CK-18 (pg/mL)
**
CD45-CD31- CD326+Zombie+ (%)
*
H
I
J
M
Ccl2 (whole lung)
*
Sftpc (whole lung)
***
Cdh1 (whole lung)
***
SYTOX+: SYTO24+
CFB-/- + H2O2
CFB+/+ + H2O2
CFB-/-+ media
CFB+/+ + media
Hours
K
CRISPR-induced CFB deletion
(3067 .. 3086) MS2131.h.CFB.F
MS2131.h.CFB.R (3480 .. 3499)
CFB
CFB mRNA
CFB
MS2131.h.CFB.sp4
MS2131.h.CFB.sp7
L
CRISPR-induced C3aR (C3aR1) deletion
C3AR1
MS571.C3aR1.g3
MS571.C3aR1.g6
MS570.C3aR1.g2
MS570.C3aR1.g16
N
SYTOX+: SYTO24+
C3aR+/+ + H2O2
C3aR-/- + H2O2
C3aR+/+ + media
C3aR-/- + media
Hours
